# Periprotein membrane lipidomics and the role of lipids in transporter function in yeast

**DOI:** 10.1101/2020.01.12.903161

**Authors:** Joury S van ‘t Klooster, Tan-Yun Cheng, Hendrik R Sikkema, Aike Jeucken, D. Branch Moody, Bert Poolman

## Abstract

The yeast plasma membrane is segregated into domains: the Micro-Compartment-of-Can1 (MCC) and Pma1 (MCP) have a different protein composition, but their lipid composition is largely unknown. We extracted proteins residing in these microdomains via stoichiometric capture of lipids and proteins in styrene-maleic-acid-lipid-particles (SMALPs). We purified SMALPs by affinity chromatography and quantitatively analyzed the lipids by mass spectrometry and their role in transporter function. We found that phospholipid and sterol concentrations are similar for MCC and MCP, but sphingolipids are enriched in MCP. Ergosterol is depleted from the periprotein lipidome, whereas phosphatidylserine is enriched relative to the bulk of the plasma membrane. Phosphatidylserine, non-bilayer lipids and ergosterol are essential for activity of Lyp1; the transporter also requires a balance of saturated/unsaturated fatty acids. We propose that proteins can function in the yeast plasma membrane by the disordered state of surrounded lipids and diffuse slowly in domains of high lipid order.

**Impact statement:** Membrane protein-specific lipidomics provides information on the organization of the yeast plasma membrane and the functioning of solute transporters

## Introduction

The interior of the cell is separated from the exterior by a lipid bilayer.(1) Cells can have more than 1,000 different lipid species, and the molecular composition at any point on a planar membrane, or in the inner and outer leaflets, differs. Eukaryotic membranes vary along the secretory pathway, and the plasma membrane is enriched in sphingolipids and sterols(2). Furthermore, the inner and outer leaflet of the plasma membrane of eukaryotic cells differ in lipid species: anionic lipids(3) and ergosterol(4) are enriched in the inner leaflet. Sphingolipids are more abundant in the outer leaflet as shown for mammalian cells. Within the membrane the lipids and proteins may cluster in domains of different composition(5).

Yeast and many other fungi tolerate a low external pH(6) and high solvent concentrations(7), which suggests a high degree of robustness of their plasma membranes. This correlates with observations that the lateral diffusion of proteins in the plasma membrane is extremely slow and the permeability for small molecules is low as compared to mammalian or bacterial membranes(8, 9), suggesting high lipid order within the plasma membrane(10). In yeast, the lateral segregation of lipids is associated with a differing location of marker proteins. Of these, the Membrane Compartments of Pma1 (MCP) and Can1 (MCC)(11, 12) are well studied. The proton-ATPase Pma1 strictly localizes to the MCP, whereas the amino acid transporters Can1 and Lyp1 localize to the MCP or MCC depending on the physiological condition(8, 13). Many more proteins are associated with MCC and MCP, but the molecular basis of their partitioning is elusive(14). The MCC is part of a larger complex called the ‘Eisosome’, which stabilizes the MCC membrane invaginations(15), leading to the term ‘MCC/eisosomes.’ In terms of lipid composition, the MCP is believed to be enriched in sphingolipids and the MCC in ergosterol(10,16,17), but evidence for this partition is indirect and mainly based on fluorophore binding(17) and lipid-dependent protein trafficking(13). Attempts to accurately determine the lipids of the yeast plasma membrane after cell fractionation are hampered by impurities from other organellar membranes.

A recently developed method involving Styrene-Maleic-Acid (SMA) polymers(18) allows extraction of proteins from their native lipid bilayer. SMA-polymers form protein-lipid containing disc-shaped structures, called ‘SMALPs’, preserve the periprotein lipids or even more distant lipid shells depending on the disc-size. We adopted this method for direct, *in situ* detection of lipids that surround named membrane proteins, which minimizes contamination of lipids from other membrane compartments. We developed a three-step method whereby very small lipid shells are generated, hereafter called the periprotein lipidomes. Shells containing a named protein are captured, and lipids associated with each named protein are detected by lipidomics in defined domains such as MCC and MCP. Finally, we used the lipidomics data to test the lipid dependence of amino acid transport by purified Lyp1 reconstituted in synthetic lipid vesicles.

## Results

### Approach to periprotein membrane lipidomics

We used SMA to extract transmembrane proteins with surrounding lipids from the plasma membrane to capture periprotein-lipid discs, called SMA-Lipid Particles SMALPs. This approach(19) avoids detergents normally needed to capture membrane proteins and selectively captures lipids within a disc of defined diameter of 9 nm ± 1 nm(18) (Fig. S1), which normally exceeds the area of most membrane proteins (∼20 nm^2^), so we expected less than 5 concentric layers of lipid. Given the density of proteins in cells of ∼ 3 per 100 nm^2^ (20) versus a SMALP area of ∼50 nm^2^ (assuming SMA-polymer contributes 1 nm to the radius), we reasoned that most SMALPs would contain a single protein and lipid would be in moderate stoichiometric excess. These predictions based on the known cross-sectional area of lipids proteins and SMALPS estimate the SMA:protein:lipid of 1:1:60-120 (Fig. 1). We refer to these lipids as the periprotein lipidome, which includes more than just the annular lipids; the latter are defined as lipids directly contacting the transmembrane domain.

**Figure 1:**
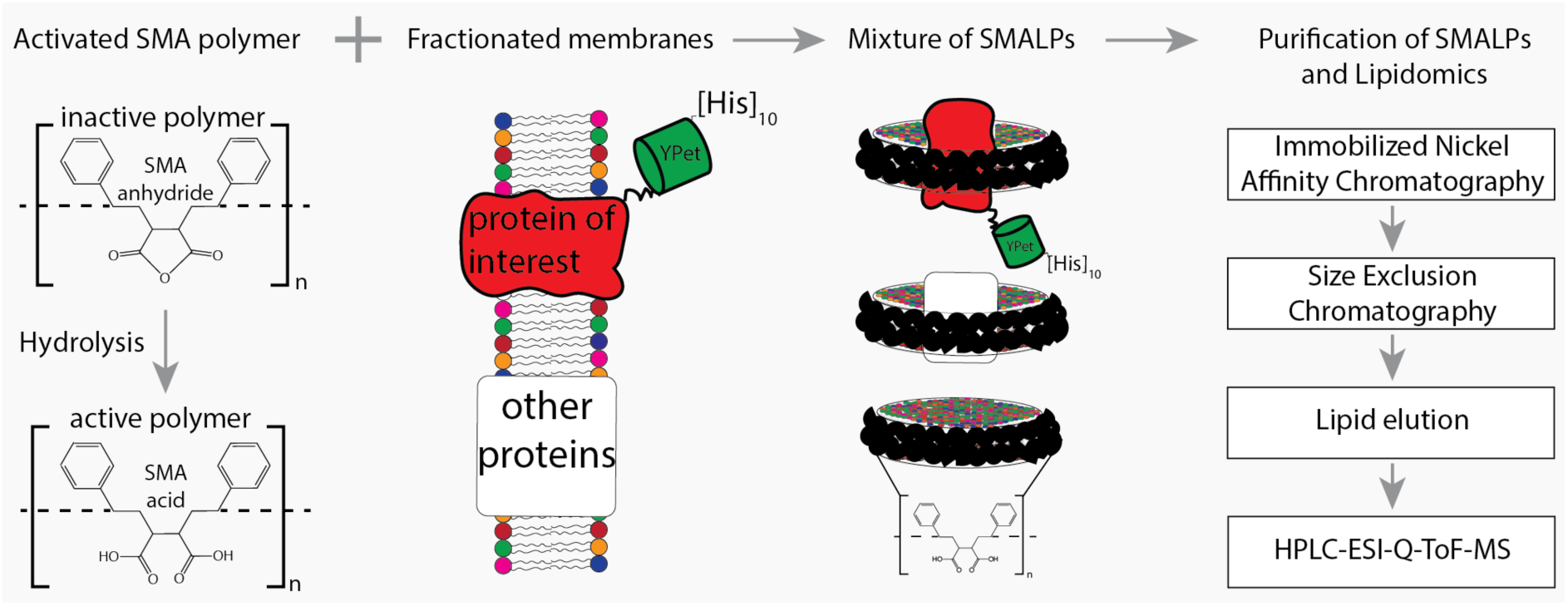
Flowchart of SMALP isolation and lipidome analysis. Hydrolysis of SMA-anhydride gives SMA acid. Combining SMA with fractionated membranes results in SMALPs in which SMA (black ribbon) has encapsulated lipids and proteins. The initial mixture consists of SMALPs containing the protein of interest (red), other proteins (white box) or no protein at all. Purification of protein-specific SMALPs is done by Immobilized Nickel Affinity Chromatography (IMAC) and Size-Exclusion Chromatography (SEC). Next, lipids associated with SMALPs are determined by reversed-phase high-performance liquid chromatography (HPLC) coupled with electrospray ionization (ESI)-quadrupole-time-of-flight (Q-ToF) mass spectrometry (MS).

We determined the lipidomes associated with Pma1, a genuine MCP resident, Sur7, a genuine MCC resident and the amino acid transporters Can1 and Lyp1, which cycle between MCP and MCC. Can1 and Lyp1 leave the MCC and are internalized from the MCP when arginine (substrate of Can1) and lysine (substrate of Lyp1) are present in excess(8, 21). At low concentrations of arginine and lysine, the Can1 and Lyp1 predominantly localize in the MCC (up to 60% of Can1 and Lyp1 molecules)(8). To trap Can1 and Lyp1 in the MCC and to obtain a better representation of protein-specific MCC lipids, we used a GFP-binding protein (GBP)(22) fused to the MCC resident Sur7 to specifically enrich for Lyp1-YPet and Can1-YPet proteins in the MCC (Fig. S2). The GFP-binding protein binds YPet with high affinity and sequesters YPet-tagged proteins, when Sur7-GBP is present in excess.

We engineered a C-terminal 10-His-tag to each of the proteins and used metal-affinity (Nickel-Sepharose) and size-exclusion chromatography for purification of SMALPs containing either Pma1-Ypet, Sur7-Ypet, Can1-YPet or Lyp1-YPet with measurable purity (Supplementary Figure 3A). SDS-PAGE analysis shows multiple protein bands (Fig. S3B), presumably due to proteolysis of protein loops during purification. Indeed, 2D native-denaturing gel electrophoresis shows that the vast majority of protein bands are genuine parts of Pma1-Ypet, Sur7-Ypet, Can1-YPet or Lyp1-YPet (Fig. S3C). Each protein migrates as a single band on a native gel and segregates into multiple bands when SDS is included in the 2^nd^ dimension of the electrophoresis. Furthermore, MS analysis of proteins in SMALPs shows peptide coverage across the full-length amino acid sequence (Fig. S3D). Finally, MS analysis of lipids extracted by SMA polymer from synthetic lipids vesicles shows that the procedure does not bias towards the extraction of specific phospholipids (Fig. S4A), similar to prior observations of sphingolipids and sterols(23, 24).

### Lipidomics analysis of periprotein microdomains

Next, the SMALP-lipid-protein complexes were treated with a chloroform-methanol-water mixture (1:2:0.8)(25) to precipitate protein and SMA polymers in the interphase and capture membrane lipids in the organic phase. Lipid mixtures were analyzed by reversed-phase high-performance liquid chromatography (HPLC) coupled with Electrospray Ionization (ESI)-Quadrupole-Time-of-Flight (QToF) mass spectrometry. Initially we generated individual lipidomes of Pma1-MCP and Sur7-MCC by comparing the lipid eluents from SMALP-protein to SMA polymer only. To detect lipid and small molecule contaminants associated with SMA, we performed negative mode comparative lipidomics(26) of SMALP-Pma1 and SMA polymer alone (Fig. 2A), which generated 815 unique ions. This documented sensitive and broad lipid detection with very low background from SMA alone, in agreement with prior analyses of conventional lipid binding proteins(27, 28).

**Figure 2.**
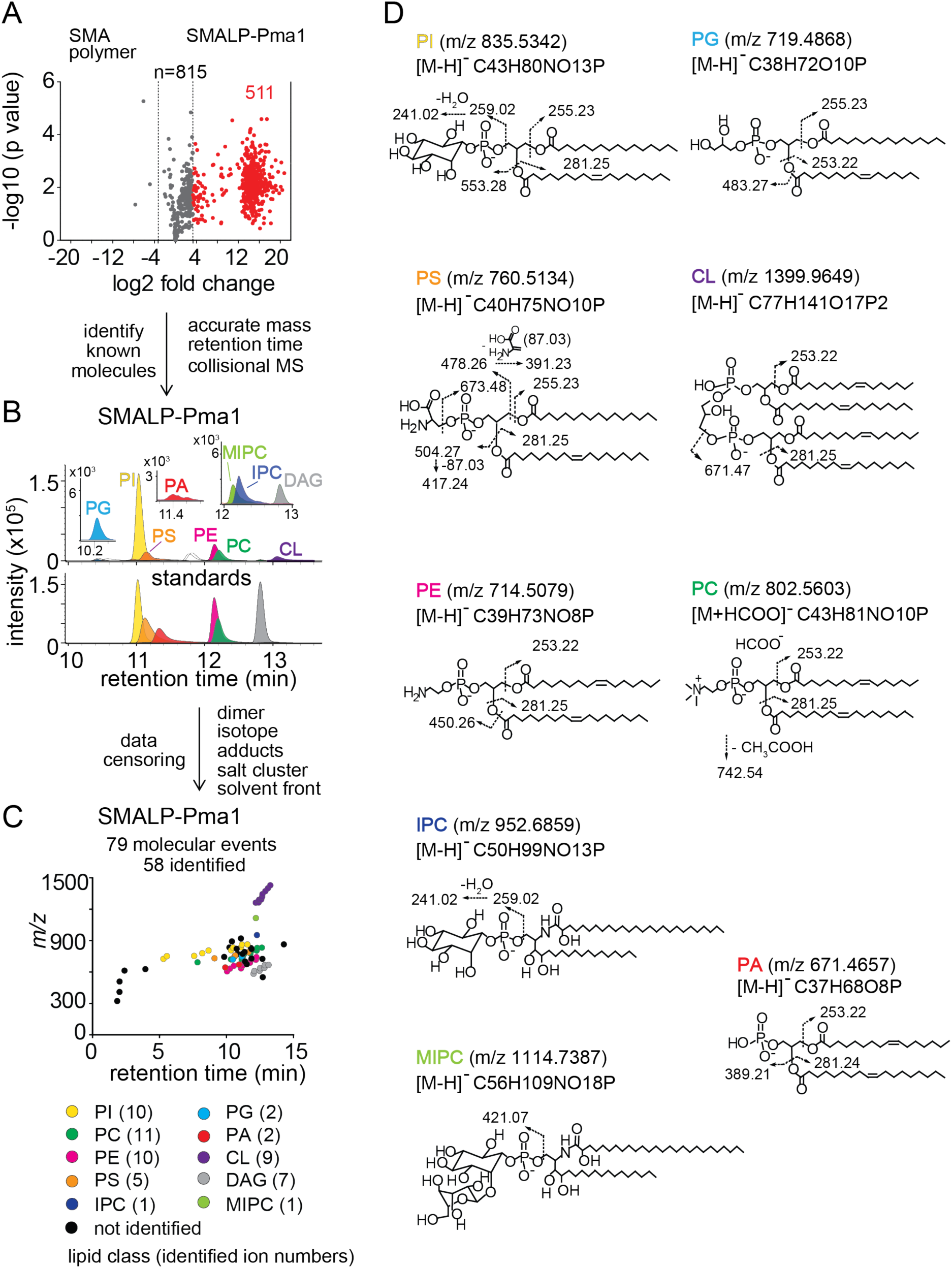
Lipidomics of SMALPs. (A) Three independently purified samples of SMALP-Pma1 obtained from yeast cells were subjected to lipid extraction, and 1 µM of the input protein was used to normalize lipid input for the reversed phase HPLC-QToF-ESI-MS in the negative ion mode. Ions were considered enriched for the SMALP-Pma1 when the intensity was at least 10-fold higher compared to the control. (B-C) Common lipids, including PI, PC, PS, PE, PG, PA, CL, DAG, IPC, and MIPC, were found based on the accurate mass, retention time or collisional MS pattern match to the synthetic standards. Chain length and unsaturation analogs were identified, whereas the redundant ions such as isotopes, lipid dimers, alternate adducts, and salt clusters were removed from the enriched ion pool. (D) Negative mode CID-MS from each lipid class identified diagnostic fragments. Molecular variants with altered chain length and unsaturation within each class were deduced based on mass intervals corresponding to CH2 or H2 hydrogens (not shown).

Next we sought to identify and remove false positive or redundant signals. Building on previously reported methods for analysis of protein-lipid complexes(28–30), we implemented data filters for lipids in protein-SMALP complexes. First, ions were considered highly Pma1-specific when signals were 10-fold higher than for SMA polymer alone. 511 ions passed this stringent criterium, whereas 3 signals were preferentially found with SMA polymer, demonstrating a low false positive lipid binding to SMALPs. We further enriched for high value hits by censoring ions present in solvents or recognizable as inorganic salt clusters based on low mass defects. We removed ions representing redundant detection of lipids as isotopes, alternate adducts and dimers [2M] (Fig. 2B). The combined effects of these filters returned 79 high quality molecular events, representing distinct retention times or m/z values.

Yet, 79 lipids still exceeded our capacity to identify them with targeted collision-induced dissociation (CID) MS, so we used a grouping technique whereby all molecules are plotted according to retention time and mass (Fig. 2C). This simple technique yields clusters comprised of lipids with the same underlying general structure with nearly co-eluting chain and unsaturation variants, represented as X:Y. Next, we solved the lead compound for key groups by CID-MS. Matching the accurate mass, retention time, and CID-MS patterns (Fig. 2B, C, and D) to standards, we identified 58 molecules in eight lipid classes, including 10 molecular species of phosphatidylinositol (PI; lead molecule, *m/z* 835.5361, 34:1), 11 phosphatidylcholines (PC; lead molecule *m/z* 804.5771, 34:1), 10 phosphatidylethanolamines (PE; m/z 716.5251, 34:1), 5 phosphatidylserines (PS; lead molecule *m/z* 760.5148, 34:1), 2 phosphatidylglycerols (PG; lead molecule *m/z* 719.4889, 32:1), 2 phosphatidic acids (PA; lead molecules *m/z* 671.4673, 34:2 PA), 9 cardiolipins (CL; lead molecule *m/z* 1399.960, 68:4), and 7 diacylglycerols (DAG; lead molecule m/z 639.5224, 34:1). Although we were unable to detect mannosyl-diinositolphosphoceramide (M(IP)_2_C), we did identify an ion matching inositolphosphoceramide (IPC, m/z 952.6846, 44:0) and mannosyl-inositolphosphoceramide (MIPC, m/z 1114.7340, 44:0) (Figure 2B). The identification of these yeast lipids was confirmed by the CID-MS (Fig. 2D).

The lipidomic profile of SMALP-Sur7 was generated by the same approach as described for SMALP-Pma1. For SMALP-Sur7, we identified 36 of 50 high quality ions, including 8 PIs, 10 PCs, 5 PEs, 7 PSs, 2 CLs, and 4 DAGs (Fig. S5) Despite the different background strains of Pma1 and Sur7, many of the same phospholipid species were found in both SMALP-protein complexes.

Given the high specificity for protein pull down and the lack of substantial adherence of lipids to SMA polymers alone, the lipids identified are likely the molecules that surround the transmembrane sequences of Pma1 and Sur7. For other SMALPs: Can1-MCP, Can1-MCC, Lyp1-MCP, and Lyp1-MCC, we carried out targeted lipidomic analysis based on the ions identified from both SMALP-Pma1 and SMALP-Sur7 lipidomes, which includes 7 classes of phospholipids, 2 types of sphingolipids, and ergosterol (supplemental table S2).

### Sphingolipids and ergosterol in MCC versus MCP domains

Ergosterol is a major component of the yeast plasma membrane. Unlike membrane phospholipids and sphingolipids that form anions, ergosterol is a neutral species that is rarely detected in negative mode lipidomics. However, in the positive mode we could detect the protonated ion [M+H]^+^ at *m/z* 397.3464, which matched the expected mass of ergosterol and had the same retention time as the ergosterol standard. MCC domains are thought to be enriched in ergosterol and sphingolipids are relatively excluded. To test this hypothesis directly, we compared the lipidomic profiles with regard to inositol-phosphoceramide (IPC), mannosyl-inositol-phosphoCeramide (MIPC) and ergosterol (Fig. 3A and S6A)). We did not initially detect IPC and MIPC in the computerized high throughput analysis of Sur7 lipidomes. These ions were missed by automated peak-picking algorithms, likely due to their very low intensity and nearby background signals. However, manual examination of ion chromatograms that focused on the mass of these two molecules in HPLC-MS convincingly detected chromatographic peaks allowing the quantitative assessment of MIPC and IPC. Unlike other lipids analyzed, standards for MIPC and IPC were not available, so we could not directly determine their molar abundance from MS signals. However, using ergosterol as an internal control, we found ∼4-fold decrease in MIPC/Ergosterol and IPC/ergosterol ratios (Fig. 3B and C) in the SMALP-Sur7-MCC compared to the SMALP-Pma1-MCP. Here, the MS signals for ergosterol are similar for Sur7-MCC and Pma1-MCP, but IPC and MIPC MS signals are different. We found similar results for Lyp1-MCP but for Can1-MCP the ratios were not significantly reduced (Fig. S6). These ratios were consistent with the conclusion that MCC-Sur7 domains have reduced sphingolipids as compared to MCP-Pma1, but inconsistent with MCC-Sur7 periprotein domains being enriched in ergosterol(16, 17). Separately, we performed ergosterol staining by filipin in our cells using Sur7-Ypet as reporter of MCC/eisosomes and did not find enhanced filipin fluorescence at the MCC domains (Fig. S7).

**Figure 3:**
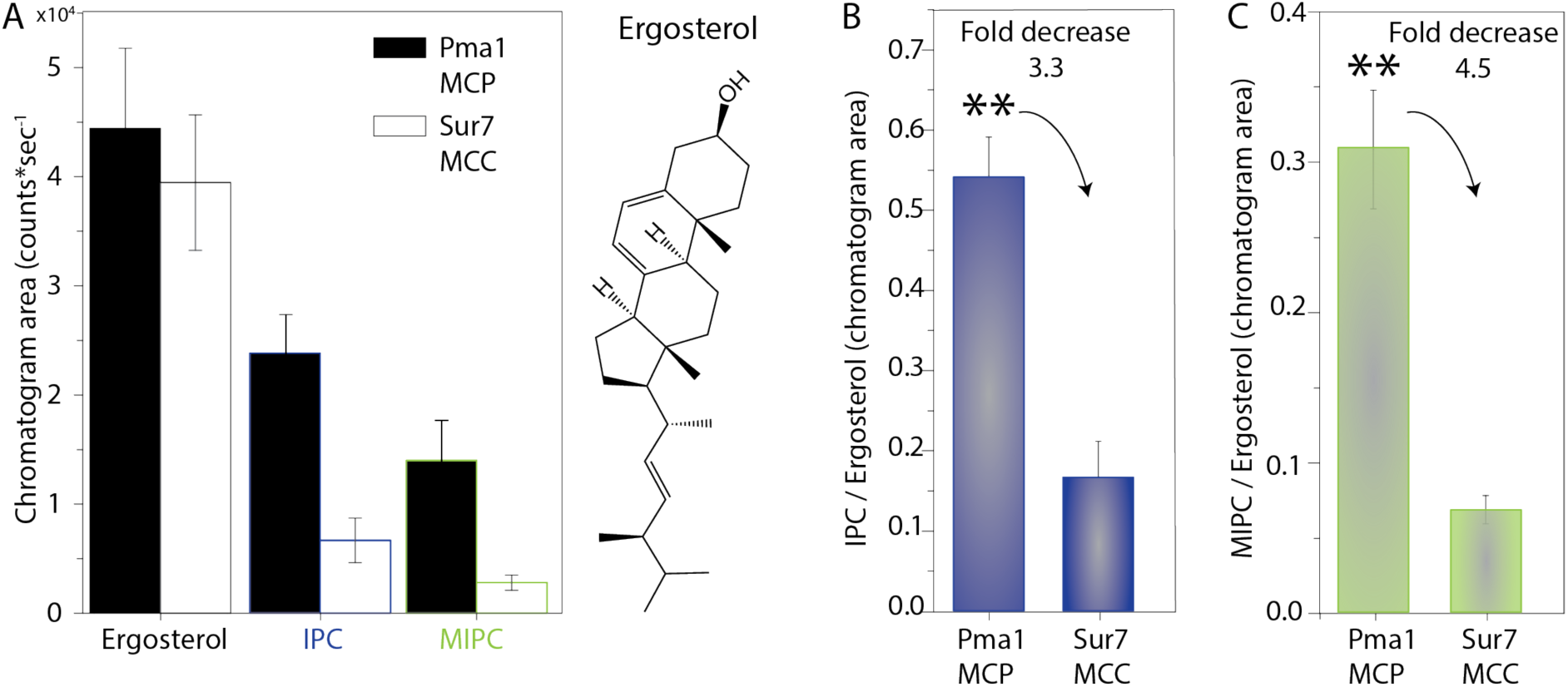
Ratios of ergosterol over sphingolipids in MCP and MCC. (A) Peak areas of ergosterol, Inositol-PhosphoCeramide (IPC) and Mannosyl-Inositol-PhosphoCeramide (MIPC) for Pma1 (MCP) and Sur7 (MCC) and structure of Ergosterol. (B) Ratio of IPC/Ergosterol for Pma1 (MCP) and Sur7 (MCC). (C) Ratio of MIPC/Ergosterol for Pma1 (MCP) and Sur7 (MCC). Number of biological replicate experiments (n) = 3, the data are presented as mean +/- standard error. P value was calculated using Student’s t-test. **,P < 0.005.

### Quantitative lipid analysis of SMALPs

From the overall lipidomes of MCC-Pma1 and MCP-Sur7, we found that many lipid species are detected in both domains. Next, we investigated whether the protein-associated domains vary in phospholipids whose enrichments and roles in these domains are unknown. We estimated the lipid quantity by comparing the peak areas of ion chromatograms to the external standard curves for PI (34:1), PC (34:1), PE (34:1), PS (34:1), PG (36:1), PA (34:1), CL (72:4), and ergosterol (Fig. S8A). Ranking the lipids by yield, we found the most abundant lipid species are PC, PI, PE, PS, and ergosterol with minor quantities of PG, PA and cardiolipin (Fig. 4A).

**Figure 4:**
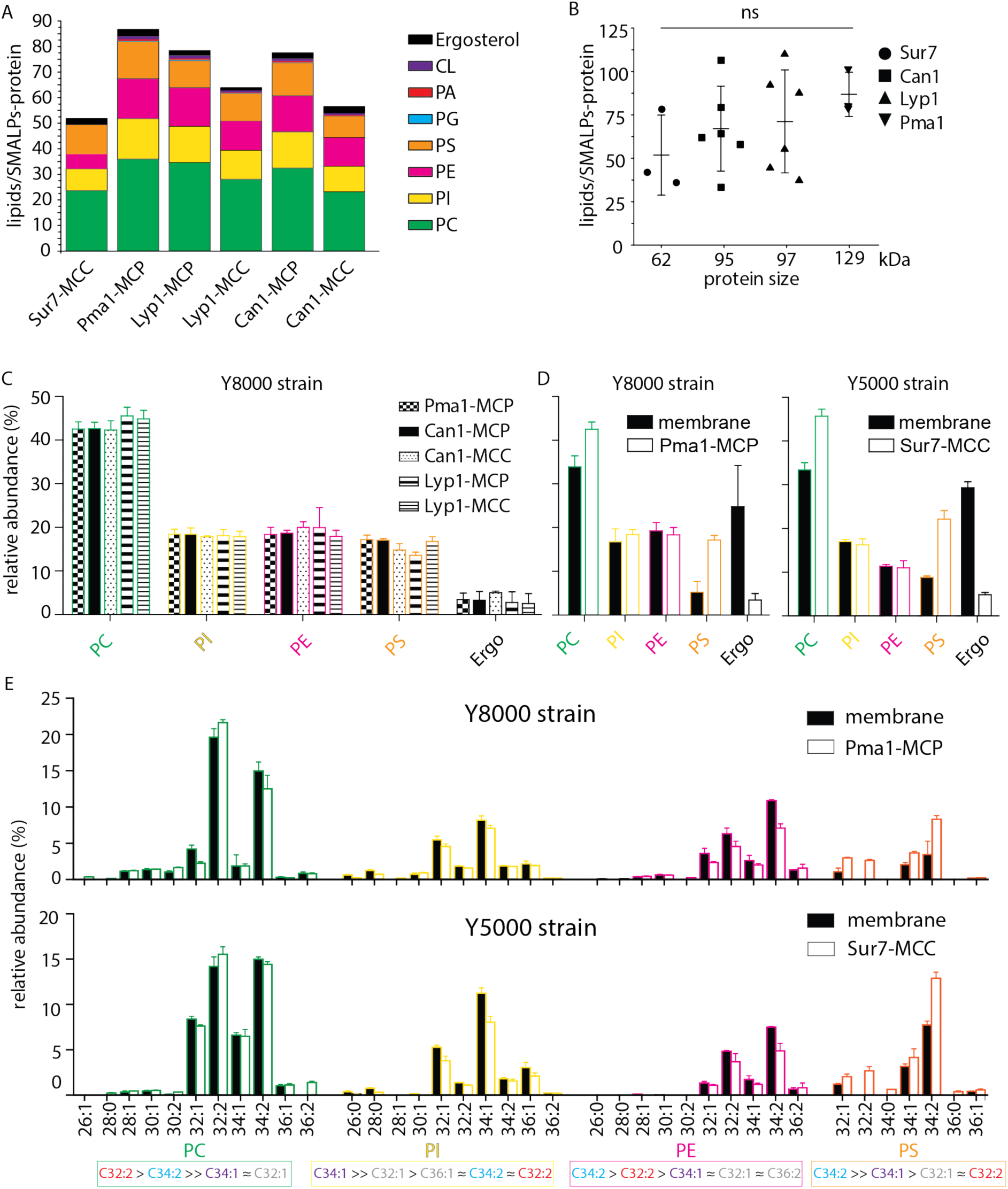
**Phospholipids of SMALPs.** (A) Number of lipid species per SMALP. (B) Total number of lipids per SMALP for each protein plotted against molecular weight of the protein, excluding the YPet moiety. (C) Relative abundance per lipid class for each SMALP. (D) Relative abundance of lipids for total plasma membrane extracts of Y8000-& Y5000-strain. (E) Phospholipid composition of the MCP and MCC. C = carbon, 1^st^ number = cumulative length of two acyl chains, 2^nd^ number = cumulative number of double bonds of two acyl chains. Phospholipid two-letter abbreviations, including color-coding: PC = PhosphatidylCholine, PE = PhosphatidylEthanolamine, PS = PhosphatidylSerine and PI = PhosphatidylInositol. Number of biological replicate experiments (n) = 3; error bars are Standard Error of the Mean (SEM).

We calculated the molar ratio of phospholipids and ergosterol associated with each SMALP-protein, assuming 1:1 stoichiometry of SMALP to protein. The estimated average numbers of phospholipids plus ergosterol per SMALP-protein complex are 52, 66, 70, and 87 associated with Sur7, Can1, Lyp1, and Pma1, respectively, which is proportional to the protein sizes (Fig. 4B). From these ratios, we estimate 1-2 rings of lipid around each protein. Since the injection amount for lipid analysis was normalized based on the same molar amount of the input protein, the smaller protein size of Sur7 could associate with less lipids and cause the fewer total ions detected in the SMALP-Sur7. However, due to variation among three independently purified SMALP-protein complexes, the protein size dependent lipids association did not reach statistical significance.

For the lipid class composition, we focused on the 4 major phospholipids and ergosterol regarding their relative abundance (Fig. 4C). To obtain better estimates of lipid quantities, we calculated conversion factors obtained from external standards and the standard addition method, which relies on true internal standards (Fig. S8 B-D). SMALPs containing Pma1, Can1, and Lyp1 were purified from the Y8000 strain, whereas SMALP-Sur7 were purified from Y5000 strain. Therefore, the total plasma membrane extracts of these two strains were also analyzed separately (Fig. 4D). We found that Y8000 samples, including Pma1-MCP, Lyp1-MCP, Lyp1-MCC, Can1-MCP, and Can1-MCC consist of similar compositions of PC (∼40%), PI (∼20%), PE (∼18%), PS (∼16%) and ergosterol (∼4%) (Fig. 4C). We also noticed that for the overall (plasma) membrane of strains Y5000 and Y8000, ergosterol (25-30 mol%) is 6-fold (Fig. 4D) higher than in the SMALPs, which suggests that ergosterol is depleted from the periprotein lipidome and more abundant in the bulk lipids surrounding the MCC and MCP proteins. Similarly, we observe 2 to 3-fold higher PS in all SMALP samples compared to overall (plasma) membrane, indicating that PS is enriched in the periprotein lipidome.

For the fatty acyl chain distribution, we found slight differences between strains. However, for both strains, we observed the similar fatty acyl chain distribution patterns in the four major phospholipid classes, as compared between the SMALP-protein associated lipids and their parent strain overall plasma membrane lipids (Fig. 4E and S9). We detected 11 forms of PC, 10 PIs, 10 PEs, and 7 PSs. We found C34:2 as major acyl chain for PE and PS and C32:2 and C34:1 for PC and PI, respectively.

In conclusion, periprotein lipidomes from SMALPs differ from the overall plasma membrane in ergosterol and PS content. Furthermore, the relative abundance of IPC and MIPC vary between the MCC and MCP, whereas differences in the major membrane phospholipids are small and not significant.

### Phospholipid dependence of transporter function

Next, we sought to validate possible functional implications of the lipidomics data by testing the lipid dependence of Lyp1, using membranes formed from synthetic lipids. We previously reconstituted Lyp1 in lipid vesicles composed of yeast total lipid extract(31). The yeast sphingolipids IPC and MIPC and certain headgroup-acyl chain combinations of the phospholipids are not available, but we could design artificial membranes that otherwise mimic those found in cells. In initial experiments with ergosterol present we observed that C34:1 (C18:1 plus C16:0 chains) palmitoyl-oleoyl-*sn*-phosphatidylX (POPX, where X=choline, ethanolamine, glycerol or serine), support a much higher transport activity than the most commonly used C36:2 forms or when C32:0 dipalmitoyl-*sn*-phosphatidylX (DPPX; C32:0 = 2 C16:0 chains) lipids (Fig. S10A) were used. When we exploit the lipidomics data and prepare vesicles with the dominant acyl chains for each lipid species using C32:2 PC and PE and C34:1 PS, which is the second most abundant PS, we also find low transport activity (Fig. S10B). Hence, we started with a mixture of POPS, POPG, POPE and POPC (17.5 mol% each) plus 30 mol% ergosterol (Fig. 5C, sample 1) and used that to benchmark the activity of Lyp1 in vesicles against the effects of phospholipids and sterols.

**Figure 5:**
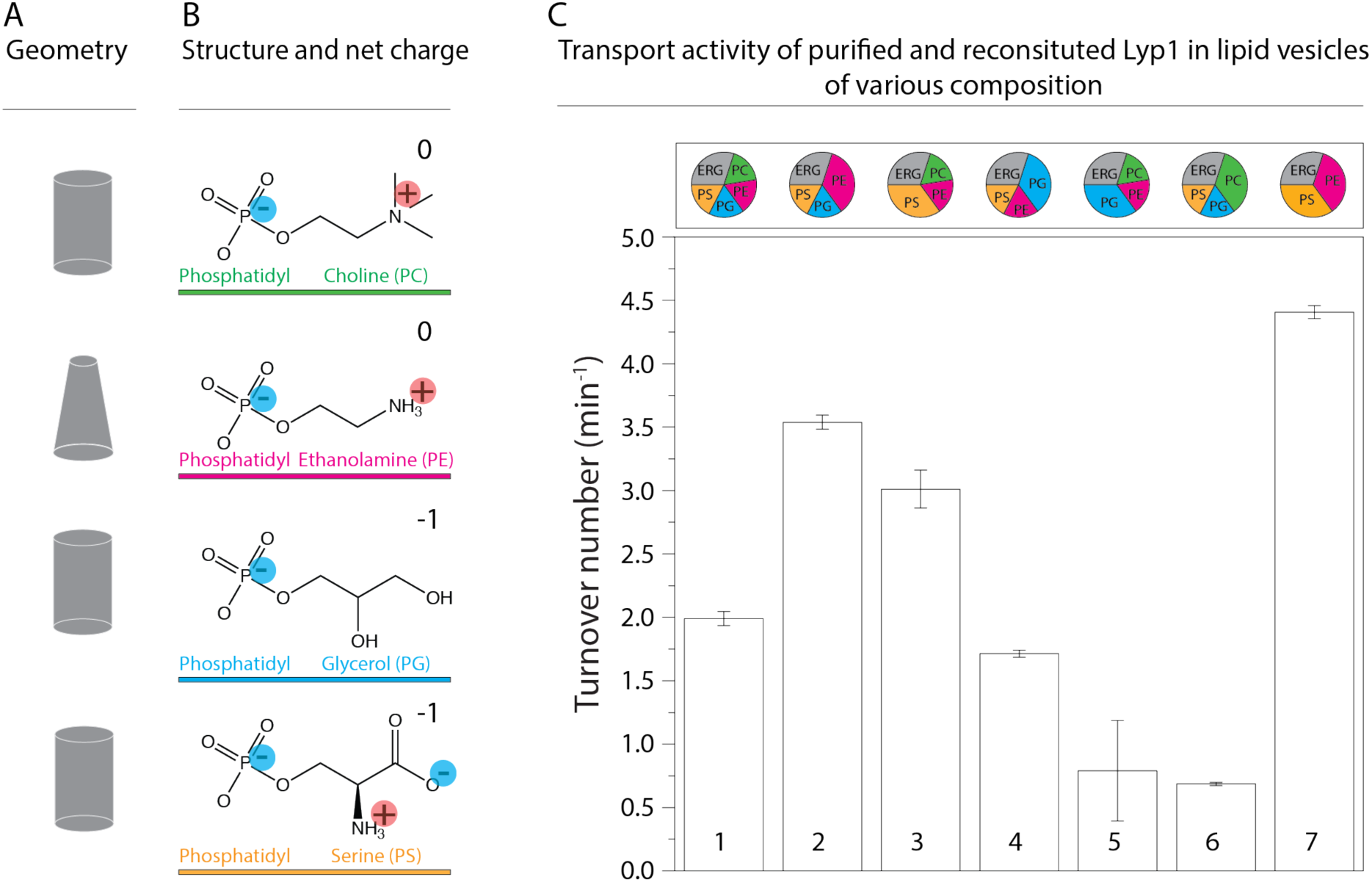
Lyp1 activity as a function of lipid composition. (A) is the geometric representation of lipids for the headgroups shown in B. (B) shows Headgroups of phospholipids and color-coding with the net charge of the lipids at neutral pH. (C) shows the turnover number of Lyp1p in different lipid mixtures with C34:1 (PO) acyl chains, where the composition (mol%) of each sample is visualized by pie-graphs, using the color coding of (B). The quantity of ergosterol was kept at 30 mol% in all mixtures. Data based on 3 replicate experiments; the bars show the standard error of the fit.

To drive the import of lysine by Lyp1, we impose a membrane potential (ΔΨ) and pH gradient (ΔpH) by diluting the vesicles containing K-acetate into Na-phosphate plus valinomycin. The magnitude of the driving force is determined by the potassium and acetate gradient (Fig. S11A). Typically, we use a ΔΨ and -ZΔpH of −81 mV each (Z=58 mV), producing a proton motive force of −162 mV. We compared the lipid composition of the Lyp1 vesicles with the starting mixture for the reconstitution, using mass spectrometry, and do not find significant differences (Fig. S4B). This rules out enrichment or depletion of certain lipids during membrane reconstitution, including the possibility that co-purified protein lipids contribute significantly to the lipid pool. Furthermore, proton permeability measurements indicate that the transport rates are not skewed by large differences in proton leakage (Fig. S12). Hence, the proton motive force is similar in vesicles of different lipid composition.

Supplementary Figure S11B shows the import of lysine over time, from which the slope estimates the initial rate of transport. Figure 5C shows that increasing either POPS or POPE increases transport activity (samples 2 and 3), while reducing the fraction of these lipids decreases the activity (samples 5 and 6) relative to that in the benchmark mixture (sample 1). Increasing the fraction of POPG or lowering of POPC (sample 4) had no negative effect on transport. This suggests that the non-bilayer lipid POPE and the anionic lipid POPS are a minimal requirement for transport (sample 7).

### The role of anionic lipids

Next, we simplified the lipid mixture by preparing vesicles composed of POPS, POPE and Ergosterol (Fig. 5C, sample 7) and step-wise reduced the quantity of PS. We found a sigmoidal relationship between Lyp1 activity and POPS concentration (Fig. 6A), which is indicative of cooperativity and suggests that more than one molecule of POPS is needed for activation of Lyp1. The anionic lipid POPG can only partly substitute for POPS (Fig. 5C, sample 5).

**Figure 6:**
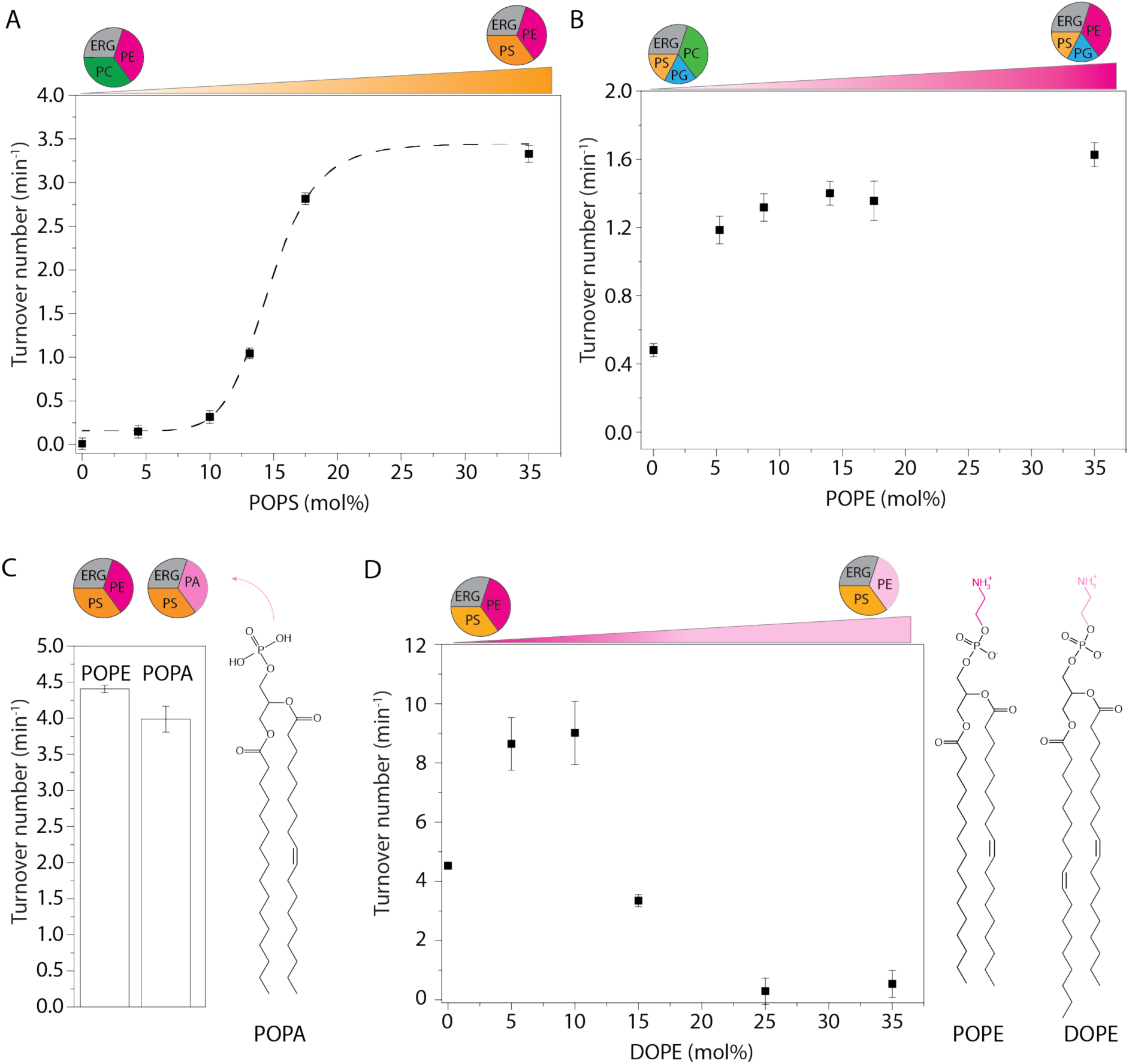
Effect of anionic and non-bilayer lipids on Lyp1 activity in proteoliposomes. (A) Turnover number of Lyp1p as a function of POPS (1-palmitoyl-2-oleoyl-*sn*-glycero-3-phosphoserine). (B) Turnover number of Lyp1 as a function of POPE (1-palmitoyl-2-oleoyl-*sn*-glycero-3-phosphoethanolamine) (mol%). (C) Turnover number of Lyp1 in vesicles with POPE versus POPA (1-palmitoyl-2-oleoyl-sn-glycero-3-phosphatidic-acid). (D) Turnover number of Lyp1 as a function of DOPE, which was increased at the expense of POPE. The triangles at the top of each graph depict the gradual replacement of one lipid for another. Number of replicate experiments (n) = 3; the bars show the standard error of the fit of n = 1; variation between replicate experiments is within 20%. Sigmoidal curves were fitted using the equation: 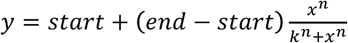

### The role of non-bilayer lipids

Many phospholipids have a cylindric geometry that allow bilayer formation with a single lipid species. The relatively small headgroup of PE, as compared to the acyl chains, results in cone geometry that does not allow bilayer formation from pure PE. PE and other non-bilayer lipids affect the lateral pressure profile and membrane curvature more drastically than bilayer forming lipids do(32). We determined the PE dependence of Lyp1 by increasing the fraction of POPE at the expense of POPC (Fig. 6B). Lyp1 transport activity increases 3-fold with increasing POPE and saturates at ∼10 mol%. To determine whether the ethanolamine headgroup or the geometric shape of the lipid is important, we substituted POPE for POPA (Fig. 6C), a non-bilayer forming conical phospholipid devoid of the headgroup moiety in PE (Fig. 5A). POPA can fully substitute POPE in transporter activity, which suggests that some lipids with non-bilayer properties are important for Lyp1 function. Next, we titrated in DOPE at the expense of POPE to gradually increase the degree of acyl chain unsaturation (Fig. 6D). We observe a 2-fold increase in Lyp1 activity with DOPE at 5 to 10 mol% and POPE at 30 to 25 mol%, but a further increase in dioleoyl at the expense of palmitoyl-oleoyl chains decreases the activity to zero. These experiments indicate that specific features of the acyl chains (or fluidity) are as important as the geometric shape of non-bilayer lipids.

### Ergosterol is essential for Lyp1 activity

Ergosterol is the major sterol of lower eukaryotes and present in the yeast plasma membrane at concentrations of ≈ 30 mol%(33), but the fraction of ergosterol in the periprotein lipid shell of Lyp1 (and Can1, Pma1 and Sur7) appears 6-fold lower than in the bulk membranes (Fig. 4B). We increased the fraction of ergosterol at the expense of POPC and observed an increase in activity up to 10 mol%. Without ergosterol, Lyp1 is not active and the activity drops above 25 mol% (Fig. 7A). As with POPS we find a sigmoidal dependence but the apparent cooperativity is much lower. Cholesterol supports less than 15% of Lyp1 activity as compared to ergosterol. Ergosterol differs from cholesterol in that it has two additional double bonds at positions C7-8 and C22-23 and one extra methyl group at C24 (Fig. 7C yellow ovals). To find out which of these is important for transport activity, we tested brassicasterol and dehydrocholesterol and an equal mixture of both (Fig. 7C). The two sterols and the mixture thereof cannot substitute for ergosterol (Fig 7B).

**Figure 7:**
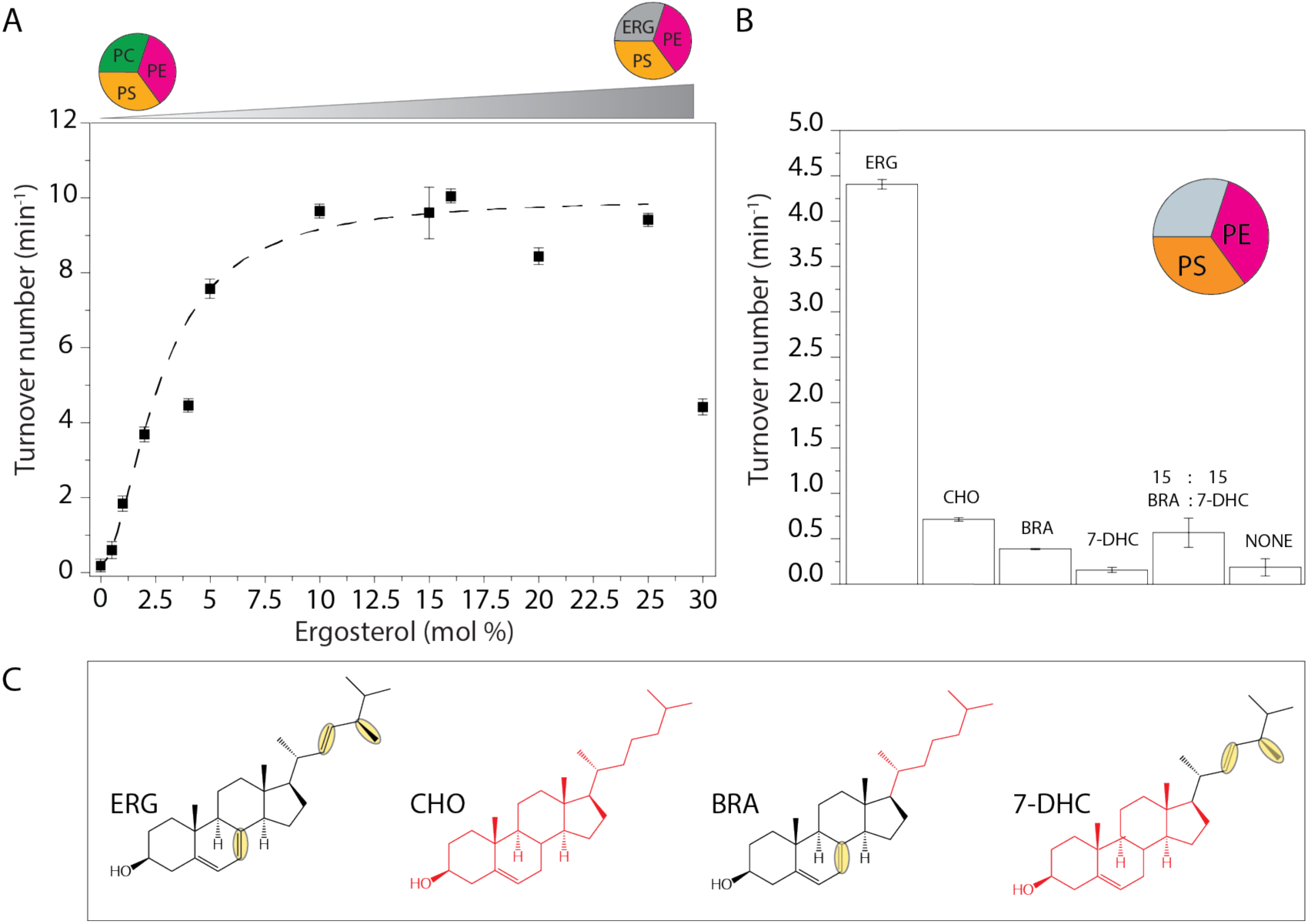
Effect of sterols on Lyp1 activity. (A) Turnover number of Lyp1 as a function of the quantity of ergosterol in the vesicles. (B) Turnover number of Lyp1 as a function of the type of sterol. (C) Structures of sterols; ERG = Ergosterol; CHO = Cholesterol; BRA = Brassicasterol; 7-DHC = 7-Dehydrocholesterol. ERG and CHO parts are black and red respectively. Structural dissimilarities between ERG and CHO are highlighted by yellow ovals. Number of biological experiments (n) = 3, variation between n = ± 20% therefor shown n = 1. The Error bars are the standard error of the fit of n = 1.

In summary, anionic lipids with saturated and unsaturated acyl chains, preferably POPS, and ergosterol are essential for Lyp1 functioning, and both lipid species stimulate transport cooperatively. This suggests that “allosteric” sites on the protein need to be occupied by specific lipids to enable transport.

### Model for protein functioning in a highly ordered yeast PM

The periprotein lipidomes and published literature lead to a new testable model (Fig. 8) of how proteins may function in a membrane of high lipid order, slow lateral diffusion and low permeability(8,9,14,34). We observe that membrane proteins like Lyp1 require a relatively high fraction of lipids with one or more unsaturated acyl chains that allow sufficient conformational flexibility of the proteins. The proteins with periprotein lipidome are embedded in an environment of lipids that are enriched in ergosterol and possibly saturated long-chain fatty acids such as present in IPC, MIPC and M(IP)_2_C), which yield a highly liquid-ordered state. The ordered state forms the basis for the robustness of the organism to survive in environments of low pH or and high solvent concentration(6, 7) and likely explains the slow lateral diffusion and the low permeability of the yeast plasma membrane (Fig. 8).

**Figure 8:**
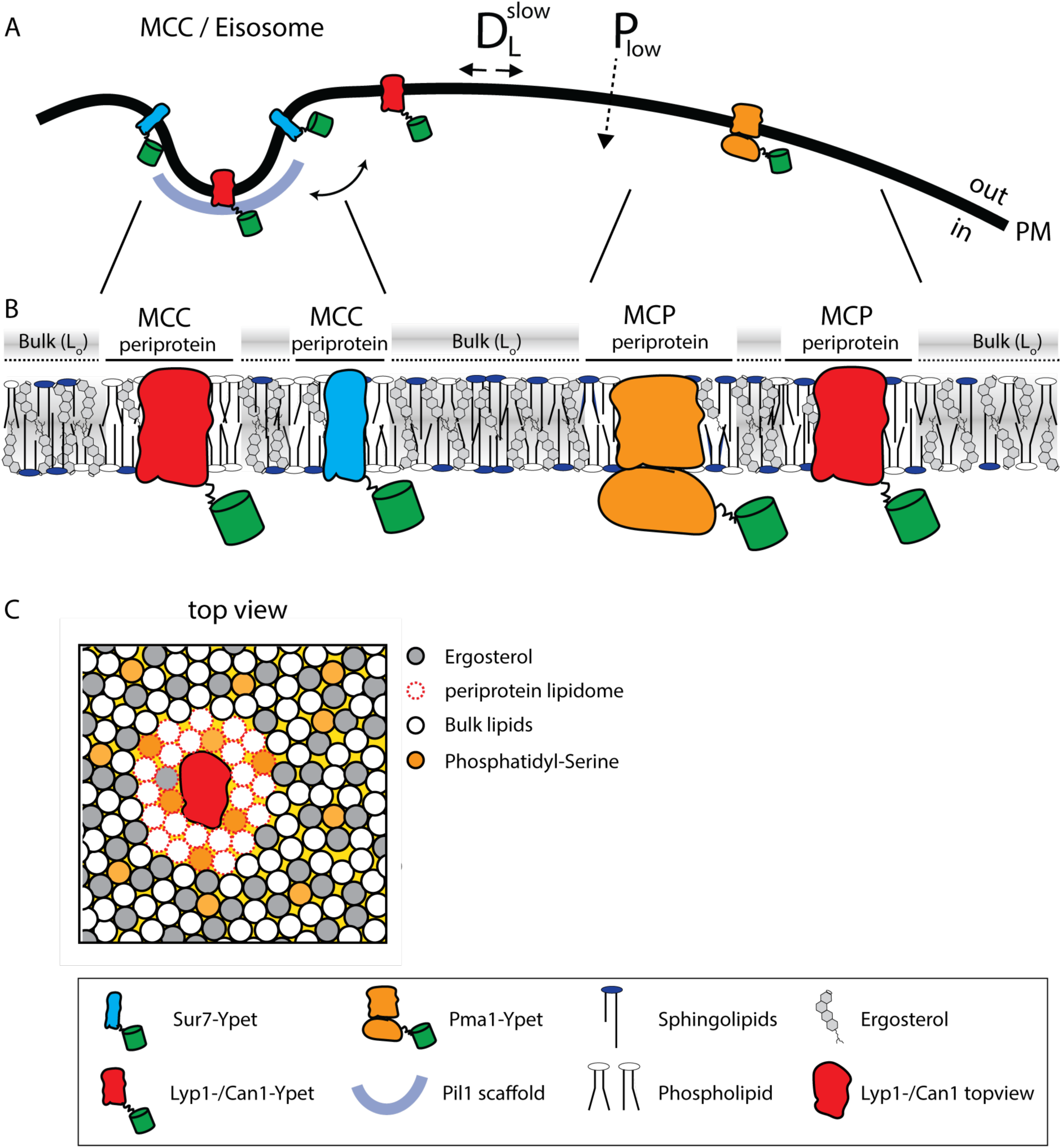
Model of the yeast plasma membrane. (A) The MCC/Eisosome is stabilized by a scaffold of Pil1 molecules. Sur7 strictly resides at the rim of the MCC/Eisosome, while Lyp1 and Can1 can diffuse in and out. Pma1 cannot enter the MCC/eisosome and resides in the MCP. (B) Schematic representation of the lipid composition of the MCC/Eisosome and the MCP based on the periprotein lipidome detected for Sur7 and Pma1, respectively. Here, phospholipids and ergosterol are similar, but sphingolipids are enriched in the MCP. The Bulk is enriched in ergosterol and represent lipids excluded from the periprotein lipidome. (C) Top view of part of the plasma membrane showing Lyp1 or Can1 enriched in phosphatidyl-serine and depleted in ergosterol as periprotein lipidome; the proteins surrounded by these periprotein lipidome diffuse very slowly in the bulk of lipids which is in a highly ordered state due to the high fraction of ergosterol. D*_L_*^slow^= slow lateral diffusion of proteins, P_low_ = low permeability of solutes, L_o_ = liquid ordered PM = plasma membrane.

How can transporters like Lyp1 function in such a membrane? The majority of transporters in the plasma membrane of yeast belong to the APC superfamily, including Lyp1 and Can1, or the Major Facilitator Superfamily (Hxt6, Gal2, Ptr2). These proteins undergo conformation changes when transiting between outward and inward conformations, which would be hindered in a highly liquid-ordered membrane, where lipids will have to be displaced when the protein cycles between the outward- and inward-facing state. To obtain an estimate of the number of displaced lipids needed for such a conformational change, we analyzed the X-ray structures of the Lyp1 homolog LeuT in different conformations(35). We analyzed structures oriented in the membrane from the OPM database(36), which positions proteins in a lipid bilayer by minimizing its transfer energy from water to membrane. We have used a numerical integration method to estimate the surface area of the outward and inward state of LeuT in the plane of the outer- and inner-leaflet at the water membrane interface (Fig. 9). We estimated the number of lipids in the vicinity of the protein by drawing an arbitrary circle around the protein. With a radius of 35 ångström from the center of the protein we need 43 to 50 lipids per leaflet depending on the conformation. For the inner-leaflet, the inward-to-outward movement of LeuT requires plus three lipids and for the outer-leaflet minus three lipids, which can probably be accommodated by local changes in membrane compressibility(37) and by redistribution of the annular and next shell of lipids even if they are surrounded by membrane that is in a highly liquid-ordered state. We find similar changes in the numbers of lipids when we analyze different conformations of membrane transporters of the MFS (not shown). Although the difference in number of lipids is small (plus or minus three), the projections in figure 9 show that the conformational changes require significant lateral displacement of lipids in both inner- and outer-leaflet. Hence, the degree of acyl chain unsaturation and low ergosterol concentration is in line with the flexibility that is needed for the conformational changes.

**Figure 9:**
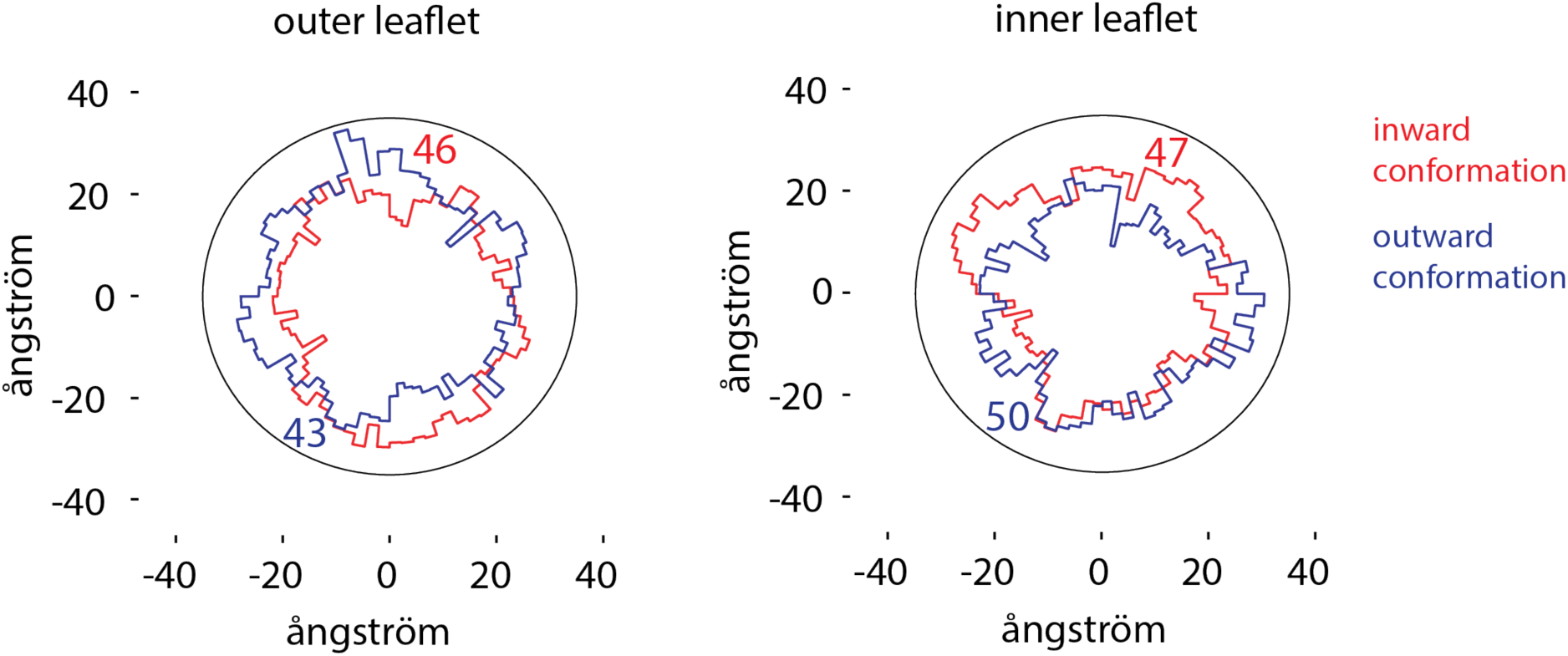
Number of lipids in inner- and outer-leaflet for the inward and outward conformation of LeuT. Protein structures were positioned in a lipid bilayer by the OPM database(36). The OPM database positions proteins in a lipid bilayer by minimizing its transfer energy from water to membrane. Shown are projections of the area occupied by inward (red), outward (purple) conformation of the protein and both projections overlaid. We have drawn a circle of 70 ångström in diameter, which represents 1-2 lipid shells based on an average area per lipid of 0.471 nm^2^(38). Finally, the number of lipids in the outer- and inner-leaflet were calculated from the difference in surface area of the circle and the protein surface.

## Discussion

Using a new method to directly quantitate individual lipids that surround named proteins, we show that the membrane proteins extracted from the MCC and MCP domains do not differ much in periprotein phospholipid and ergosterol composition, but do differ significantly in sphingolipid content. Furthermore, we find a much lower fraction of ergosterol and an enrichment of PS in the lipidome associated with Can1, Lyp1, Pma1 and Sur7 than in the surrounding bulk lipids. Assuming one protein per SMALP, we find between 52 to 87 lipids per protein. The number of lipids increases when sphingolipids are included and agrees with estimates reported in the literature(18). Calculated from the estimated perimeter of Lyp1 homolog LeuT, the number of lipids corresponds to 1-2 shells of lipids surrounding the protein in the SMALP. Hence, we conclude that the SMA extraction targets the periprotein lipidome of membrane proteins. Our *in vitro* analysis of the activity of the amino acid transporter Lyp1 reveals the requirements of the protein for specific lipids, and the optimal conditions require a stringent degree of acyl chain saturation, and amounts of anionic lipids (>15 mol%), non-bilayer lipids (>10 mol%) and ergosterol (5 mol%), which match the observed lipids in the HPCL-MS analysis. Except for phosphatidylserine we do not find a strong effect of a specific lipid headgroup on the activity of Lyp1.

When we benchmark our *S. cerevisiae* lipidomic data to other studies of the yeast plasma membrane we find similar, small quantities of PA, CL and PG(39–42). Thus, the majority of protein is extracted from the plasma membrane rather than from internal membranes. In terms of selectivity, we benefit from the ability to trap Can1 and Lyp1 in MCC/eisosomes, taking advantage of the GFP-binding protein(22) and the fact that we purify proteins using an affinity tag. Thus, our approach for lipid analysis is much less hampered by contamination with internal membranes than conventional fractionation studies. The SMALP technology only identifies protein specific lipidomes and does not report the overall composition of the plasma membrane.

Reported values for the average degree of acyl chain unsaturation for the plasma membrane of yeast vary. The values we find in our SMALPs are consistent with(41), but somewhat higher than found by others using non-SMALP methods(39,40,43,44). We detected the sphingolipids IPC and MIPC and expected to also find M(IP)_2_C as major lipid species(41), but our M(IP)_2_C signal detection was poor in the overall (plasma) membrane. We therefore do not provide specific conclusions about the quantities of M(IP)_2_C in our SMALP samples. We find similar amounts of ergosterol in MCC and MCP by mass spectrometry analysis, whereas filipin staining in the literature has suggested that ergosterol is enriched in MCC/eisosomes(45). Here both the lipidomics data and filipin staining are consistent with one another(46), but not with prior observations(16, 17).

We find that 30 mol% of ergosterol, corresponding to the overall concentration of this molecule in the plasma membrane, reduces the activity of Lyp1 (Fig. 7A). The observed 3-5 mol% of ergosterol in SMALP-Lyp1 is more in line with a high transport activity. Detailed analysis of the effects of lipids on the activity and regulation of membrane transport is limited to studies on bacterial transporters such as the lactose-proton symporter LacY(47), a leucine-proton symporter(48), the ATP-driven betaine transporter OpuA(49) and other membrane-associated proteins(50, 51). Each of these systems requires anionic (PG) and non-bilayer (PE) lipids, and function optimally in lipids with dioleoyl chains. We find that Lyp1 performs poorly in dioleoyl-*sn*-phosphatidyl-based lipids and has an order of magnitude higher activity in palmitoyl-oleoyl-*sn*-phosphatidyl lipids.

Estimates of global membrane order, using fluorescence lifetime decay measurements and mutants defective in sphingolipid or ergosterol synthesis, suggest that the yeast plasma membrane harbors highly-ordered domains enriched in sphingholipids(10, 52). This conclusion is consistent with the observation that the lateral diffusion of membrane proteins in the plasma membrane of yeast is 3-orders of magnitude slower than has been observed for membranes in the liquid-disordered state(8, 13). Accordingly, the passive permeability of the yeast plasma membrane for weak acids is orders of magnitude lower than in bacteria(9). A membrane in the gel or highly liquid-ordered state may provide low leakiness that is needed for fungi to strive in environments of low pH and/or high alcohol concentration, but it is not compatible with the dynamics of known membrane transporters, which undergo relatively large conformational changes when transiting from an outward- to inward-facing conformation. In fact, we are not aware of any transporter that is functional when it is embedded in a membrane in the liquid-ordered state.

The large, ordered lipid domains observed previously(10) might represent the MCP because Pma1 is enriched for sphingolipids relative to Sur7 in MCC. Furthermore, ergosterol is depleted from the periprotein lipidome of the Sur7, Can1, Lyp1 and Pma1 proteins and is 6-fold enriched in the surrounding bulk membranes. Sterols are known to increase the lipid order and interact more strongly with saturated than unsaturated lipids, therefore depletion of ergosterol from the periprotein lipidome is expected to decrease the lipid order. We propose that large parts of the yeast plasma membrane are in the liquid-ordered state but that individual proteins are surrounded by one or two layers of lipids with at least one unsaturated acyl chain to enable sufficient conformational dynamics of the proteins, which might be needed for transporters like Lyp1(35,53,54). The proteins in these membrane domains would diffuse slowly because they are embedded in an environment with a high lipid order.

## Methods

### Yeast strains and plasmids

*Saccharomyces cerevisiae* strains (Table S1) are derived from Σ1278b (from Bruno Andre(55)) or BY4709; all strains are uracil auxotrophs (Ura3-negative). Plasmids are based on pFB001(31) and constructed by User Cloning™(New England Biolabs) in a three-way ligation method, where the plasmid is amplified in three similar sized fragments of which one contains the Gene Of Interest (GOI) and the other two form the full amp^R^ gene when ligation is successful. Plasmid fragments were transformed and ligated in *Escherichia coli* MC1061 by means of the heat shock procedure. Plasmid assembly and nucleotide sequences were confirmed by DNA sequencing, and plasmids isolated from *E. coli* were transformed to *Saccharomyces cerevisiae* using the Li-acetate method(56) and selection was based on Ura3 complementation. Positive transformants were re-cultured twice to ensure clonality.

### Cultivation of S. cerevisiae and protein expression

Chemicals were purchased from Sigma-Aldrich (DE), unless otherwise indicated. Cells were cultured at 30°C with 200 RPM shaking in 50 mL CELLreactor™ filter top tubes (*Greiner Bio-On)*. Strains were grown overnight in 5 mL synthetic maltose media lacking uracil, lysine, arginine and histidine. Lack of uracil ensures the plasmid is retained, while lack of lysine, arginine and histidine ensures stable expression of Lyp1 and Can1 in the yeast PM. Media was prepared by dissolving (2% w/v maltose), 0.69% yeast nitrogen base (YNB) without amino acids. Necessary amino acids were supplemented using 0.16% Kaiser synthetic mixture without uracil, lysine, arginine and histidine(57) *i.e. a mixture containing specific amino acids* except *uracil, lysine, arginine and histidine* (Formedium, UK). After growth overnight, cultures were diluted to OD_600_ = 0.15 in 50 mL of Synthetic Maltose media lacking ura, lys, arg and his and grown to OD_600_ = 0.75-1.5 in a 250 mL Erlenmeyer flask. After ∼ 8 h, cultures were diluted 50-fold into 1.6L synthetic maltose media lacking ura, lys, arg and his; after ∼15 h, the OD_600_ reached 1.0-2.0, after which 1% *w/v* galactose was added to induce protein expression for 2, 2.5 and 3 hours for Can1, Lyp1 and Pma1, respectively. To validate protein expression, we co-expressed the fusion constructs YPet-POI and mCherry-Pil1 and used confocal laser scanning microscopy (*Zeiss LSM710*); mCherry-Pil1 is a reporter of MCC/eisosomes. The excitation lasers for Ypet and mCherry were set at 488 and 543nm with emission filter settings between 509-538nm and 550-700nm, respectively; the pinhole was set to 1μm. Images were processed using ImageJ to adjust contrast and brightness levels. (Original files are accessible in the supplementary data.) The expression of Lyp1 in *Pichia pastoris* and its membrane isolation was described previously(31).

### Plasma membrane isolation

*P. pastoris* and *S. cerevisiae* cultures were harvested by centrifugation at 7,500 x g, for 15 min at 4°C (Beckman centrifuge J-20-XP, Rotor JLA 9.1000, US). Further steps were performed at 4°C or on ice. The resulting cell pellet was resuspended in 150 mL of cell resuspension buffer (CRB; 20mM Tris-HCl pH6.7, 1mM EGTA, 0.6M sorbitol, 10μM pepstatin A (Apollo Scientific, UK), 10μM E-64 (Apollo Scientific, UK) plus protease inhibitors (cOmplete Mini EDTA-free™, ROCHE, 1 tablet/75 mL). Centrifugation was repeated and the resulting cell pellet was resuspended to OD_600_ = 200 in CRB with 1 tablet cOmplete Mini EDTA-free™/10 mL). The resulting cell suspension was broken using a cell disrupter (Constant cell disruption systems, US) by three sequential passes operating at 39Kpsi, after which 2 mM fresh PMSF was added. Unbroken cells were pelleted by centrifugation at 18,000 RCF for 30 min at 4°C (Beckman centrifuge J-20-XP, Rotor JA16.250, DE). The supernatant was transferred to 45 TI rotor (Beckman, DE) tubes and membranes were pelleted by ultracentrifugation (Optima XE-90, Beckman, DE) at 186,000 RCF, 90 min at 4°C. The resulting membrane pellet was resuspended to homogeneity at 400 mg/mL using a potter Elvehjem tissue grinder in 20 mM Tris-HCl pH7.5, 0.3M Sucrose, 10μM Pepstatin A, 10μM E-64 (Apollo Scientific, UK), 2 mM PMSF and protease inhibitor cocktail (cOmplete Mini EDTA-free™, 1 tablet/10 mL). Aliquots of 2 mL were snap frozen using liquid nitrogen and stored at −80°C until further use.

### SMA preparation

Styrene Maleic Acid anhydride polymer (Xiran-SZ30010, Polyscope, NL) was hydrolyzed as described(58) and freeze-dried in aliquots of 2 grams. Upon use, SMA polymer was dissolved using 50 mM Tris-HCl pH7.5 at 0.1 g/mL.

### SMALP formation and protein purification

Styrene Maleic Acid Lipid Particles (SMALPs) were formed by combining *S. cerevisiae* membranes with SMA polymer at a ratio of 1:3 w/w and left for 16 hours at 30°C. Unextracted material was removed by ultracentrifugation at 186,000 x g 60 min at 4°C. Further steps were performed at 4°C or on ice. The supernatant was mixed with 1 mL Ni-Sepharose resin (GE healthcare, US) and incubated for 24 hours under gentle nutation, and then poured into an empty column. The Ni-Sepharose was washed with 10mL Tris-HCl pH7.5, and SMALPs were eluted in 3 sequential steps using 1mL 50 mM Tris-HCl, 50 mM Imidazole pH7.5 with 10 minutes incubation between each elution. All elution fractions were pooled and concentrated to 500 μL, using a 100 kDa spin concentrator (Merck, DE). Next, the sample was applied onto a size-exclusion chromatography column (Superdex 200 increase 10/300GL, GE Healthcare, US) attached to an HPLC (Åkta, Amersham bioscience, SE) with in-line fluorescence detector (1260 Infinity, Agilent technologies, US) (FSEC) set to excitation and emission wavelengths of 517 and 530 nm, respectively, with bandwidths of 20 nm. 500 μL Fractions were collected and concentrated to 20-40 μL. The SMALP concentration was determined from absorbance measurements at 517 nm using a nandrop (ND-1000, Isogen lifescience, NL) and an extinction coefficient for YPet of 26.810 M*cm^-1^. Samples were flash frozen in liquid nitrogen and stored at −80°C until further use.

### Lipid standards

Diacylglycerol (DAG, #800515) was purchased from Sigma-Aldrich. Phosphatidylcholine (PC 34:1, #850475), phosphatidylinositol (PI 34:1, #850142), phosphatidylserine (PS 34:1, #840032), phosphatidylethanolamine (PE 34:1, #850757), phosphatidylglycerol (PG 36:1, #840503), phosphatidic acid (PA 34:1, #840857), cardiolipin (CL 72:4, #710335) were purchased from Avanti polar lipids.

### Lipid extraction and mass spectrometry

Lipid extraction from the SMALPs and the crude membranes was performed based on the Bligh and Dyer method(25). The lower organic phase was separated from the upper aqueous phase and dried under a nitrogen stream. The extracted lipid residue was re-dissolved in the starting mobile phase A and the injection volume was 10 µl for each HPLC-MS run. The injection concentration for lipid extracts from SMALPs was normalized to 1 µM based on input protein concentration. The injection amount for SMA polymer control was 0.02 µg. The injection amounts for the crude membranes were 0.05, 0.25, 1, 2.5, or 10 µg depending on the application. The samples were run on an Agilent Poroshell 120 A, EC-C18, 3 x 50 mm, 1.9 µm reversed phase column equipped with an Agilent EC-C18, 3 x 5 mm, 2.7 µm guard column and analyzed using Agilent 6530 Accurate-Mass Q-ToF/ 1260 series HPLC instrument. The mobile phases were (A) 2 mM ammonium-formate in methanol /water (95/5; V/V) and (B) 3 mM ammonium formate in 1-propanol/cyclohexane/water (90/10/0.1; v/v/v). In a 20-minute run, the solvent gradient changes as follows: 0-4 min, 100% A; 4-10 min, from 100% A to 100% B; 10-15 min, 100%B; 15-16 min, from 100% B to 100% A; 16-20 min, 100% A. For the lipidomic analysis, three independently purified SMALP-Pma1 (MCP) or SMALP-Sur7 (MCC) complexes were analyzed and compared to the SMA polymers alone. Data were analyzed using Mass Hunter (Agilent) and R package XCMS(59) for lipidomic peak analyses and *in house* designed software methods(26).

For the mass spectrometry belonging to figure S4: 10ul of the lipid extraction was injected on a hydrophilic interaction liquid chromatography (HILIC) column (2.6 μm HILIC 100 Å, 50 × 4.6 mm, Phenomenex, Torrance, CA). The mobile phases were (A) acetonitrile/acetone (9:1, v/v), 0.1% formic acid and (B) acetonitrile/H2O (7:3, v/v), 10mM ammonium formate, 0.1% formic acid. In a 6.5-minute run, the solvent gradient changes as follows: 0-1 min, from 100% A to 50% A and B; 1-3 min, stay 50% B; 3-3.1 min, 100% B; 3.1-4 min, stay 100% B; 4-6.5 min from 100% B to 100% A. Flowrate was 1.0mL/min The column outlet of the LC system (Accela/Surveyor; Thermo) was connected to a heated electrospray ionization (HESI) source of mass spectrometer (LTQ Orbitrap XL; Thermo) operated in negative mode. Capillary temperature was set to 350°C, and the ionization voltage to 4.0 kV. High resolution spectra were collected with the Orbitrap from m/z 350-1750 at resolution 60,000 (1.15 scans/second). After conversion to mzXML data were analyzed using XCMS version 3.6.1 running under R version 3.6.1.

To estimate the lipid quantity, the lipid concentration was obtained by external standard curve fitting. Briefly, a series of concentrations of synthetic standard were prepared and analyzed by HPLC-MS to determine the response factor and degree of linearity of input lipid to count values. The ion chromatogram peak areas from known concentrations were used to generate the standard curves for determining the unknown concentrations of the extracted lipids. For key applications where the highest quality lipid quantification was needed or when input lipid mixtures were complex, we used internal standards for authentic lipids and the method of standard addition. For phospholipids, 0.25 µg crude membrane samples were spiked with a series of known concentrations (0, 0.5, 1.0, 1.5, and 2.0 pmol/µl) of synthetic molecules for C34:1 PC, C34:1 PE, C34:1 PS, or C34:1 PI and subjected to HPLC-MS negative ion mode analysis. For ergosterol, 0.05 µg crude membrane samples were spiked with a series of known concentrations of the synthetic standard (0, 1 pmol/µl, 3 pmol/µl, and 5 pmol/µl), and the data were acquired in the positive ion mode. The ion chromatogram peak areas of the specific *m/z* values were plotted against the concentrations of the spiked synthetic standards to extrapolate concentration of natural lipids on the X-axis.

### Liposome formation

Phospholipids were obtained from (Avanti polar lipids Inc, Alabaster, AL, USA), and brassicasterol (CarboSynth, UK), 7-dehydrocholesterol (Sigma Aldrich, DE), cholesterol and ergosterol (Sigma Aldrich, DE) were obtained from the indicated vendor. Lipids were dissolved in chloroform and mixed at desired ratios (mol%) to a total weight of 10 mg in a 5 mL glass round bottom flask. Chloroform was removed by evaporation at 40°C and an applied pressure (*p*) of 370 mBar, using a rotary vaporizer (rotavapor r-3-BUCHI). The resulting lipid film was resuspended in 1 mL of diethylether and the previous step was repeated at atmospheric pressure until a dry film was visible. To remove any residual solvent a pressure of 10 mBar was applied for 20 minutes. The obtained lipid film was hydrated in 1mL of 50 mM NH_4_Acetate pH7.5, 50 mM NaCl by shaking for 5 min and then transferred to a plastic tube compatible with sonication. The lipid suspension was homogenized by tip sonication with a Sonics Vibra Cell sonicator (Sonics & Materials Inc.) at amplitude of 70% for 2 minutes with 5 sec pulses and 5 sec pauses between each pulse. The sample was kept at 4°C in ethanol:water:ice (25:25:50 v/v/v). The lipid suspension (1 mL aliquot at 10 mg of lipid/mL) was transferred to a 1.5 mL Eppendorf tube, snap frozen in liquid nitrogen and thawed at 40 °C. This process was repeated 4 times and stored in liquid nitrogen until further use.

### Proteoliposome preparation

Protein purification and membrane reconstitution was performed as described(31). Briefly, *n*-Dodecylmaltoside (DDM) was used to solubilize Lyp1-GFP. The protein was purified by Immobilized Metal Affinity Chromatography (IMAC, using Nickel-Sepharose) and Fluorescence Size-Exclusion Chromatography (FSEC, using a Superdex 200 increase 300/10 GL column attached to an Åkta 900 chromatography system (Amersham Bioscience, SE) with in-line Fluorescence detector (1260 Infinity, Agilent technologies, US). In parallel, liposomes were thawed at room temperature and subsequently homogenized by 11 extrusions through a 400nm polycarbonate filter (Avestin Europe GMBH, Ger). Next, liposomes were destabilized using Triton X-100 and titrated to a point beyond the saturation point (Rsat) as described(60); the final turbidity at 540 nm was at approximately 60% of Rsat. Purified Lyp1 was mixed with triton x-100-destabilized liposomes of the appropriate lipid composition at a protein-to-lipid ratio of 1:400 and incubated for 15 min under slow agitation at 4°C. Bio-beads SM-200 (Biorad, Hercules, Ca, USA), 100 mg/0.4% Triton X-100 (final concentration of Triton x-100 after solubilization of the liposomes) were sequentially added at 15, 30 and 60 min. The final mixture was incubated overnight, after which a final batch of bio-beads was added and incubation continued for another 2 hours. Protein containing liposomes (proteo-liposomes) were separated from the Bio-beads by filtration on a column followed by ultracentrifugation 444,000 x g at 4°C for 35 min. Proteo-liposomes were suspended in 10 mM potassium-phosphate plus 100 mM potassium-acetate pH 6.0, snap frozen and stored in liquid nitrogen.

### In vitro transport assays

Transport assays were performed and the formation of a proton gradient (ΔpH) and membrane potential (**ΔΨ**) was established as described in(31). Both ΔpH and **ΔΨ** were formed by diluting the proteo-liposomes (suspended in 10 mM potassium-phosphate plus 100 mM potassium-acetate pH 6.0) 25-fold into 110 mM sodium-phosphate pH6.0 plus 0.5 µM valinomycin. This results in maximal values of ZΔpH and **ΔΨ** of −83 mV as calculated according to the Nernst equation, yielding a proton motive force of −166 mV.

### Data analysis and transport rates

The data were analyzed in Origin (OriginLab, MA). The initial rates of transport were determined from the slope of progress curves (e.g. Supplementary Figure S11B). The relative transport activity was determined by normalizing each vesicle preparation to the sample with a lipid composition POPC:POPE:POPS:POPG:Ergosterol in a ratio of 17.5:17.5:17.5:17.5:30 (mol%).

### Protein surface area calculations

Membrane-oriented protein structures were obtained from the OPM database(36), the OPM database positions proteins in a lipid bilayer by minimizing its transfer energy from water to membrane; LeuT inward-open (PDB ID: 3f3a), LeuT outward-open (PDB ID: 5jag). Protein area in the plane of the outer and inner leaflet determined by the OPM database were calculated by numerical integration as follows: equation: 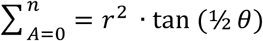. For this a polar coordinate system from the center of the protein was established (n=80) resulting in an angle (θ) of 4.5° between each coordinate. The distance (r) was determined from the most distant atom between two polar coordinates. This distance was applied at ½ θ. From distance r on ½ θ, a perpendicular line was drawn (b) resulting in a right-angled triangle for which the surface area can be calculated using the tangent, ½ θ and r. The area was calculated for each of the 80 polar coordinates and summated to acquire the total surface area of the protein.

## Acknowledgements

This work was carried out within the BE-Basic R&D Program, which was granted a FES subsidy from the Dutch Ministry of Economic affairs, agriculture and innovation (EL&I), and was supported by an ERC Advanced Grant (ABCVolume; #670578) and by NIH grants (AI116604 and AR048632). The research was also funded by NWO TOP-PUNT (project number 13.006) grants. We thank Dr. prof. Bruno André for sharing the 23344C and SG067 strains.

## Author contributions

J.S.v.t.K., D.B.M. and B.P designed the research plan; J.S.v.t.K performed the research except for the liquid-chromatography coupled to mass spectrometry analysis; T-Y.C performed the LC-MS analysis; J.S.v.t.K., T-Y.C., D.B.M. and B.P. analyzed the data; H.R.Sikkema performed the *in silico* analysis of protein surface calculations; A. Jeucken performed mass spectrometry analysis corresponding to figure S4; J.S.v.t.K., T-Y.C., D.B.M. and B.P. wrote the paper.

## Source data

Source data is uploaded separately as excel files:

Lipidomics SMALP Can1-MCC

Lipidomics SMALP Can1-MCP

Lipidomics SMALP Lyp1-MCC

Lipidomics SMALP Lyp1-MCP

Lipidomics SMALP Pma1-MCP

Lipidomics SMALP Sur7-MCC

Lipidomics Y5000

Lipidomics Y8000

## SUPPLEMENTARY INFORMATION

### Supplementary tables

**Table S1:** All strains and plasmids used in this study

| Strain/Name | Genotype | Ref / Origin |
| --- | --- | --- |
| Y8001/Lyp1 MCP | *ura3-Δ1*, pJK2001 | (<sup>13</sup>) 23344C |
| Y8002/Can1 MCP | *ura3-Δ1*, pJK2002 | (<sup>13</sup>) 23344C |
| Y8003/Pma1 MCP | *ura3-Δ1*, pJK2003 | (<sup>13</sup>) 23344C |
| Y10001/Lyp1 MCC | Sur7p-GBP, Pil1-mCherry, *can1-Δ1*, *gap1-Δ1*, *ura3-Δ1*, pJK2001 | (<sup>13</sup>) SG067 |
| Y10002/Can1 MCC | Sur7p-GBP, Pil1-mCherry, *can1-Δ1*, *gap1-Δ1*, *ura3-Δ1*, pJK2002 | (<sup>13</sup>) SG067 |
| Y5007/Sur7 | *ura3-Δ1*, Sur7p-Ypet | This Study |
| Plasmid | Description | origin |
| pJK2001 | CEN-ARS, pGal_Lyp1p-Ypet_RGShis10, *ura3* | This study |
| pJK2002 | CEN-ARS, pGal_Can1p-Ypet_RGShis10, *ura3* | This study |
| pJK2003 | CEN-ARS, pGal_Pma1p-Ypet_RGShis10, *ura3* | This study |

**Table S2:** lipids identified by targeted lipidomic analysis based on the ions identified from both SMALP-Pma1 and SMALP-Sur7. Other SMALPs: Lyp1-MCP, Lyp1-MCC, Can1-MCP, and Can1-MCC. Values are expressed as pmol lipid per 10 pmol protein. Chromatography peak areas used in our calculations are given for ergosterol and the sphingolipids IPC and MIPC.

| m/z/lipid chain | pmol lipid / 10 pmol protein |  |  |  |  |  |
| --- | --- | --- | --- | --- | --- | --- |
|  | Pma1 MCP | Sur7 MCC | Lyp1 MCP | Lyp1 MCC | Can1 MCP | Can1 MCC |
| **PC** |  |  |  |  |  |  |
| 692.4508/C26:1 | 3.03 ± 0.13 | 0 ± 0 | 2,56 ± 0,49 | 2,45 ± 0,34 | 2.33 ± 0.37 | 1.42 ± 0.32 |
| 722.4977/C28:0 | 1.11 ± 0.13 | 1.06 ± 0.24 | 1,01 ± 0,23 | 0,98 ± 0,2 | 0.98 ± 0.17 | 0.7 ± 0.09 |
| 720.4821/C28:1 | 9.93 ± 0.8 | 2.23 ± 0.47 | 10,52 ± 2,61 | 8,57 ± 2,27 | 8.13 ± 1.47 | 6 ± 0.81 |
| 748.5134/C30:1 | 11.75 ± 1.13 | 2.42 ± 0.5 | 9,19 ± 2,6 | 9,74 ± 2,43 | 10.48 ± 1.82 | 7.41 ± 1.11 |
| 746.4972/C30:2 | 13.38 ± 1.14 | 1.71 ± 0.57 | 10,18 ± 2,08 | 11,01 ± 2,42 | 10.79 ± 1.71 | 8.07 ± 1.11 |
| 776.5447/C32:1 | 18.11 ± 0.69 | 37.82 ± 10.28 | 24,89 ± 9,15 | 16,54 ± 3,04 | 18.09 ± 2.32 | 13.81 ± 1.41 |
| 774.529/C32:2 | 177.7 ± 17.5 | 76.5 ± 19.7 | 160,6 ± 455 | 134,0 ± 32,6 | 159.9 ± 38.9 | 112.5 ± 26.4 |
| 804.576/C34:1 | 15.21 ± 1.45 | 31.44 ± 7.41 | 13,52 ± 4,48 | 13,79 ± 4,38 | 12.76 ± 2.21 | 9.22 ± 2 |
| 802.5603/C34:2 | 101.06 ± 3.49 | 70.8 ± 17.72 | 99,03 ± 28,42 | 76,83 ± 10,1 | 92.97 ± 14.04 | 66.37 ± 14.55 |
| 832.6073/C36:1 | 1.98 ± 0.18 | 5.68 ± 2 | 3 ± 1,17 | 1,63 ± 0,19 | 2.22 ± 0.79 | 1.35 ± 0.46 |
| 830.5916/C36:2 | 6.62 ± 1.27 | 7.01 ± 2.29 | 11,88 ± 6,78 | 4,96 ± 1,68 | 6.15 ± 1.99 | 5.28 ± 2.26 |
| **PI** |  |  |  |  |  |  |
| 725.4246/C26:0 | 1.98 ± 0.28 | 0.44 ± 0.14 | 1,65 ± 0,46 | 1,81 ± 0,46 | 1.97 ± 0.56 | 1.12 ± 0.28 |
| 753.4559/C28:0 | 5.92 ± 0.71 | 1.47 ± 0.38 | 4,58 ± 1,13 | 4,57 ± 1,24 | 5.25 ± 1.32 | 3.31 ± 0.69 |
| 751.4403/C28:1 | 1.4 ± 0.15 | 0 ± 0 | 0,8 ± 0,2 | 1,01 ± 0,24 | 0.97 ± 0.24 | 0.61 ± 0.16 |
| 779.4716/C30:1 | 7.48 ± 1.04 | 0 ± 0 | 4,49 ± 1,02 | 5,75 ± 1,73 | 5.09 ± 1.66 | 4.03 ± 1 |
| 807.5029/C32:1 | 37.64 ± 4.87 | 18.31 ± 4.05 | 34,09 ± 9,66 | 28,14 ± 8,04 | 34.62 ± 7.99 | 23.3 ± 5.7 |
| 805.4872/C32:2 | 13.04 ± 1.24 | 5.23 ± 1.23 | 10,31 ± 2,33 | 9,22 ± 2,01 | 10.09 ± 1.75 | 7.35 ± 1.64 |
| 835.5342/C34:1 | 58.15 ± 7.45 | 40.37 ± 11.86 | 55,76 ± 18,5 | 41,09 ± 10,71 | 55.14 ± 13.58 | 38.64 ± 9.53 |
| 833.5185/C34:2 | 14.73 ± 1.53 | 7.9 ± 2.61 | 13,1 ± 4,06 | 10,33 ± 2,51 | 13.11 ± 3.06 | 9.28 ± 2.19 |
| 863.5655/C36:1 | 15.74 ± 1.54 | 10.89 ± 3.98 | 15,29 ± 5,58 | 10,96 ± 2,52 | 14.44 ± 3.27 | 10.8 ± 2.35 |
| 861.5498/C36:2 | 1.37 ± 0.06 | 0.99 ± 0.31 | 1,26 ± 0,45 | 0,9 ± 0,2 | 1.18 ± 0.24 | 0.92 ± 0.19 |
| **PE** |  |  |  |  |  |  |
| 606.414/C26:0 | 0.65 ± 0.01 | 0 ± 0 | 0 ± 0 | 0 ± 0 | 0 ± 0 | 0 ± 0 |
| 634.4453/C28:0 | 0.73 ± 0.02 | 0 ± 0 | 0,56 ± 0,13 | 0,49 ± 0,12 | 0.54 ± 0.06 | 0.47 ± 0.07 |
| 632.4296/C28:1 | 3.57 ± 0.4 | 0 ± 0 | 2,6 ± 0,61 | 2,77 ± 0,89 | 3.11 ± 0.42 | 2.23 ± 0.49 |
| 660.4609/C30:1 | 4.9 ± 0.47 | 0 ± 0 | 3,06 ± 0,69 | 3,62 ± 1,06 | 4 ± 0.43 | 3.03 ± 0.55 |
| 658.4453/C30:2 | 2.2 ± 0.4 | 0 ± 0 | 1,17 ± 0,28 | 1,61 ± 0,57 | 1.9 ± 0.26 | 1.35 ± 0.34 |
| 688.4922/C32:1 | 19.14 ± 1.23 | 5.41 ± 1.61 | 16,34 ± 3,8 | 13,46 ± 3,04 | 14.88 ± 1.84 | 12.48 ± 2.44 |
| 686.4766/C32:2 | 37.88 ± 6.66 | 17.38 ± 3.42 | 29,59 ± 6,36 | 29,36 ± 8,28 | 35.51 ± 4.26 | 26.77 ± 6.61 |
| 716.5235/C34:1 | 16.34 ± 1.65 | 5.97 ± 1.94 | 13 ± 4,72 | 10,18 ± 2,83 | 10.59 ± 1.5 | 10.15 ± 1.23 |
| 714.5079/C34:2 | 58.31 ± 7.03 | 23.42 ± 5.13 | 50,17 ± 12,15 | 43,34 ± 9,29 | 57.73 ± 8.83 | 43.95 ± 10.36 |
| 742.5392/C36:2 | 13.33 ± 4.01 | 4.74 ± 2.92 | 35 ± 27,9 | 10,07 ± 5,34 | 12.35 ± 5.41 | 11.95 ± 7.86 |
| **PS** |  |  |  |  |  |  |
| 732.4821/C32:1 | 24.41 ± 2.05 | 9.66 ± 2.03 | 15,08 ± 4,08 | 16,61 ± 4,51 | 18.06 ± 2.52 | 12.68 ± 3 |
| 730.4664/C32:2 | 21.59 ± 2.16 | 12.63 ± 2.03 | 16,25 ± 4,36 | 18,05 ± 5,27 | 21.47 ± 4.31 | 13.24 ± 4.21 |
| 760.5134/C34:1 | 29.86 ± 1.69 | 21.4 ± 7.29 | 19,22 ± 6,22 | 19,11 ± 4,33 | 21.8 ± 2.91 | 15.24 ± 3.5 |
| 758.4977/C34:2 | 68.04 ± 6.34 | 63.41 ± 16.1 | 52,99 ± 15,47 | 51,66 ± 13,49 | 67.27 ± 12.28 | 43.25 ± 12.91 |
| 788.5447/C36:1 | 1.9 ± 0.08 | 2.69 ± 0.43 | 1,9 ± 0,6 | 1,24 ± 0,12 | 0 ± 0 | 0 ± 0 |
| **PG** |  |  |  |  |  |  |
| 719.4868/C32:1 | 0.84 ± 0.33 | 0 ± 0 | 1.73 ± 0.79 | 0.71 ± 0.5 | 0.72 ± 0.42 | 0.18 ± 0.06 |
| 747.5181/C34:1 | 0.95 ± 0.31 | 0 ± 0 | 1.73 ± 0.77 | 0.75 ± 0.47 | 0.82 ± 0.45 | 0.3 ± 0.12 |
| **PA** |  |  |  |  |  |  |
| 643.4344/C32:2 | 3.24 ± 0.45 | 0 ± 0 | 2.04 ± 0.87 | 1.96 ± 0.79 | 2.72 ± 0.91 | 1.67 ± 0.7 |
| 671.4657/C34:2 | 3.2 ± 0.28 | 0 ± 0 | 3.09 ± 1.37 | 1.9 ± 0.63 | 2.83 ± 0.83 | 2 ± 0.7 |
| **CL** |  |  |  |  |  |  |
| 1261.8241/C58:3 | 0.62 ± 0.05 | 0 ± 0 | 0.5 ± 0.18 | 0.42 ± 0.12 | 0.5 ± 0.09 | 0.33 ± 0.05 |
| 1263.8397/C58:2 | 0.54 ± 0.03 | 0 ± 0 | 0 ± 0 | 0 ± 0 | 0.31 ± 0.02 | 0.26 ± 0.06 |
| 1289.8554/C60:3 | 1.35 ± 0.12 | 0 ± 0 | 1.24 ± 0.45 | 0.89 ± 0.3 | 1.07 ± 0.27 | 0.76 ± 0.17 |
| 1291.871/C60:2 | 0.84 ± 0.16 | 0 ± 0 | 1.08 ± 0.38 | 0.5 ± 0.04 | 0.85 ± 0.18 | 0.44 ± 0.1 |
| 1317.8867/C62:3 | 1.22 ± 0.12 | 0 ± 0 | 1.08 ± 0.4 | 0.78 ± 0.26 | 0.94 ± 0.26 | 0.65 ± 0.16 |
| 1343.9023/C64:4 | 1.39 ± 0.14 | 0 ± 0 | 1.3 ± 0.48 | 0.89 ± 0.32 | 1.1 ± 0.28 | 0.72 ± 0.17 |
| 1371.9336/C66:4 | 2.56 ± 0.27 | 0 ± 0 | 2.7 ± 1 | 1.73 ± 0.58 | 2.12 ± 0.57 | 1.46 ± 0.37 |
| 1399.9649/C68:4 | 2.33 ± 0.23 | 0.5 ± 0.21 | 2.39 ± 0.92 | 1.56 ± 0.48 | 1.92 ± 0.58 | 1.33 ± 0.33 |
| 1427.9962/C70:4 | 0.88 ± 0.1 | 0 ± 0 | 0.89 ± 0.32 | 0.59 ± 0.15 | 0.72 ± 0.23 | 0.52 ± 0.13 |
| **ergosterol** |  |  |  |  |  |  |
| 397.3459 | 28.17 ± 5.61 | 24.45 ± 4.61 | 11.95 ± 6.15 | 20 ± 12.93 | 22.54 ± 5.98 | 27.65 ± 6.06 |
| **Chromatography peak areas** |  |  |  |  |  |  |
| **ergosterol** |  |  |  |  |  |  |
| 397.3459 | 44390 ± 7358 | 39465 ± 6204 | 31142 ± 18607 | 21443 ± 9362 | 36755 ± 8084 | 43658 ± 8191 |
| **IPC** |  |  |  |  |  |  |
| 952.6846/C42:0 | 23809 ± 3506 | 6679 ± 2040 | 21481 ± 7033 | 12324 ± 5434 | 9134 ± 2505 | 10023 ± 2452 |
| **MIPC** |  |  |  |  |  |  |
| 1114.735/C42:0 | 14005 ± 3680 | 2797 ± 697 | 16561 ± 6280 | 6231 ± 2545 | 5020 ± 2244 | 4784 ± 944 |

### Supplementary figures

**Figure S1.**
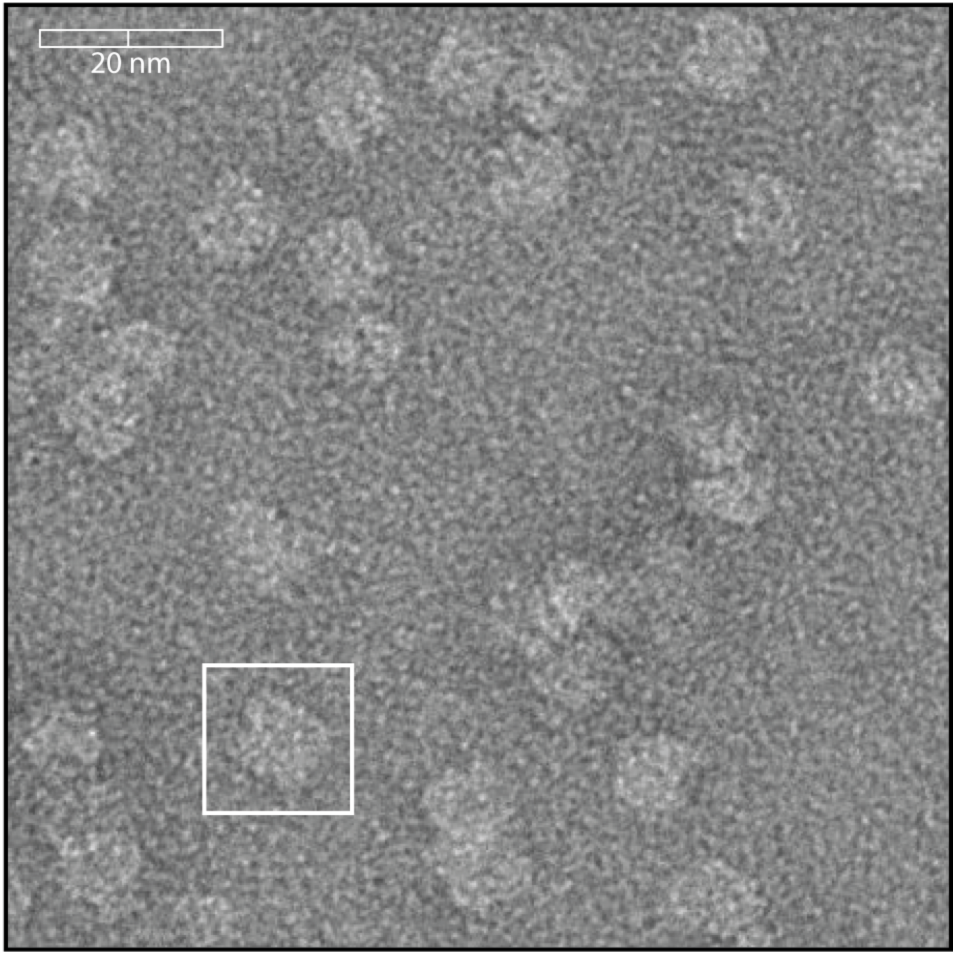
Negative stain Cryo-EM of SMALPs. SMALPs containing Lyp1-Ypet were visualized by negative stain cryo-EM. Scale bar is indicated on top left. The white box highlights one SMALP, which is approximately 10 nm.

**Figure S2:**
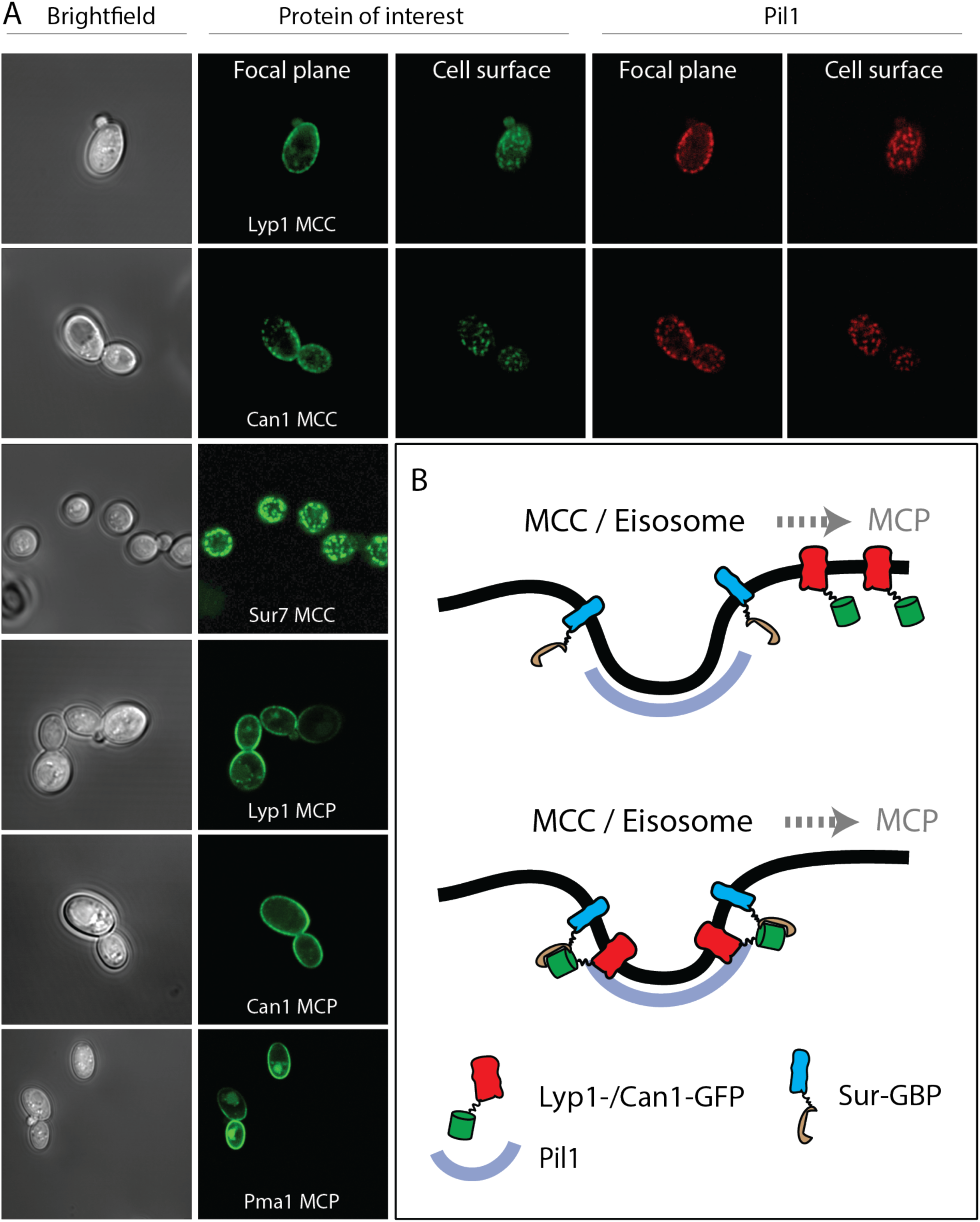
Microscopy images of strains used in this study. (A) Brightfield images depict *S. cerevisiae* visualized with visible light. (B) Proteins of interest are visualized by a Green Fluorescent Protein (Ypet) fused to Lyp1, Can1, Sur7 or Pma1. MCC = Micro Compartment of Can1; MCP = Micro Compartment of Pma1. Eisosomal protein Pil1 is visualized by a Red Fluorescent Protein (mCherry) and co-localizes with proteins in the MCC. B. Membrane protein trapping mechanism as designed by Rothbauer et al^22^ and used by^13^. GFP binding protein (GBP) is fused to Sur7, and Lyp1 and Can1 are fused to GFP. Upon interaction between GBP and GFP, Can1 and Lyp1 are trapped in the MCC/Eisosome.

**Figure S3:**
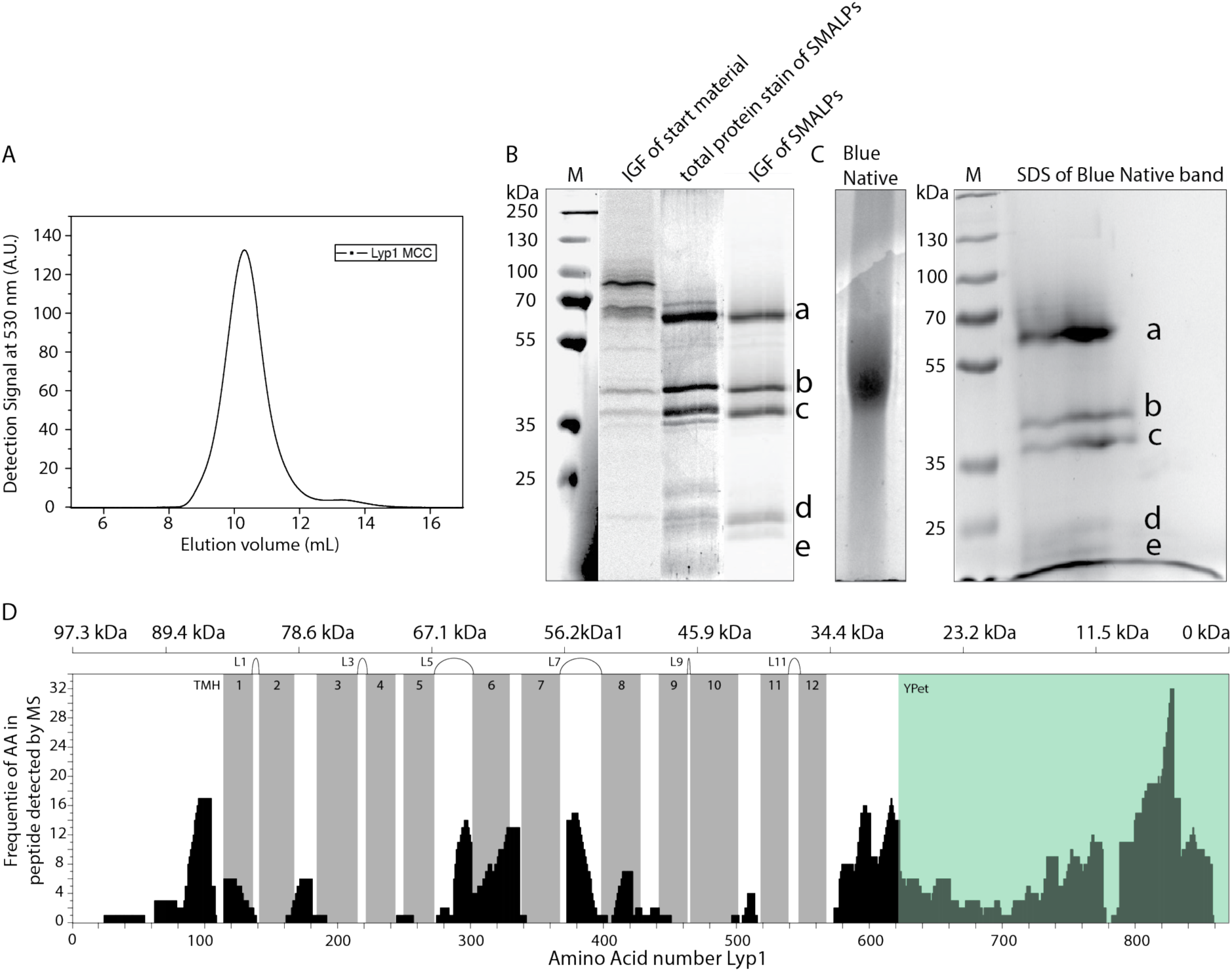
Polyacrylamide gel electrophoresis and mass spectrometry analysis of Lyp1-Ypet SMALPs. (A) Fluorescence Size-Exclusion Chromatography profile of Lyp1-Ypet SMALPs on a Superdex 200/30 GL column. (B) SDS-PAGE and (C) Blue Native-PAGE of Lyp1Ypet-SMALPs. M = Marker, IGF = In Gel Fluorescence of Ypet. Corresponding bands in SDS-PAGE and 2D native-denaturing gel electrophoresis are indicated by a-e. (D) Peptide coverage of Lyp1Ypet-SMALPs measured by Mass Spectrometry. TMH = Trans Membrane Helix, Gray bars indicate the position of each TMH. Extracellular loops are indicated as Lx, where x = the loop number. The top scale indicates the molecular weight of randomized protein fractions in kDa (kiloDalton) starting from the C-terminal YPet, highlighted in green.

**Figure S4:**
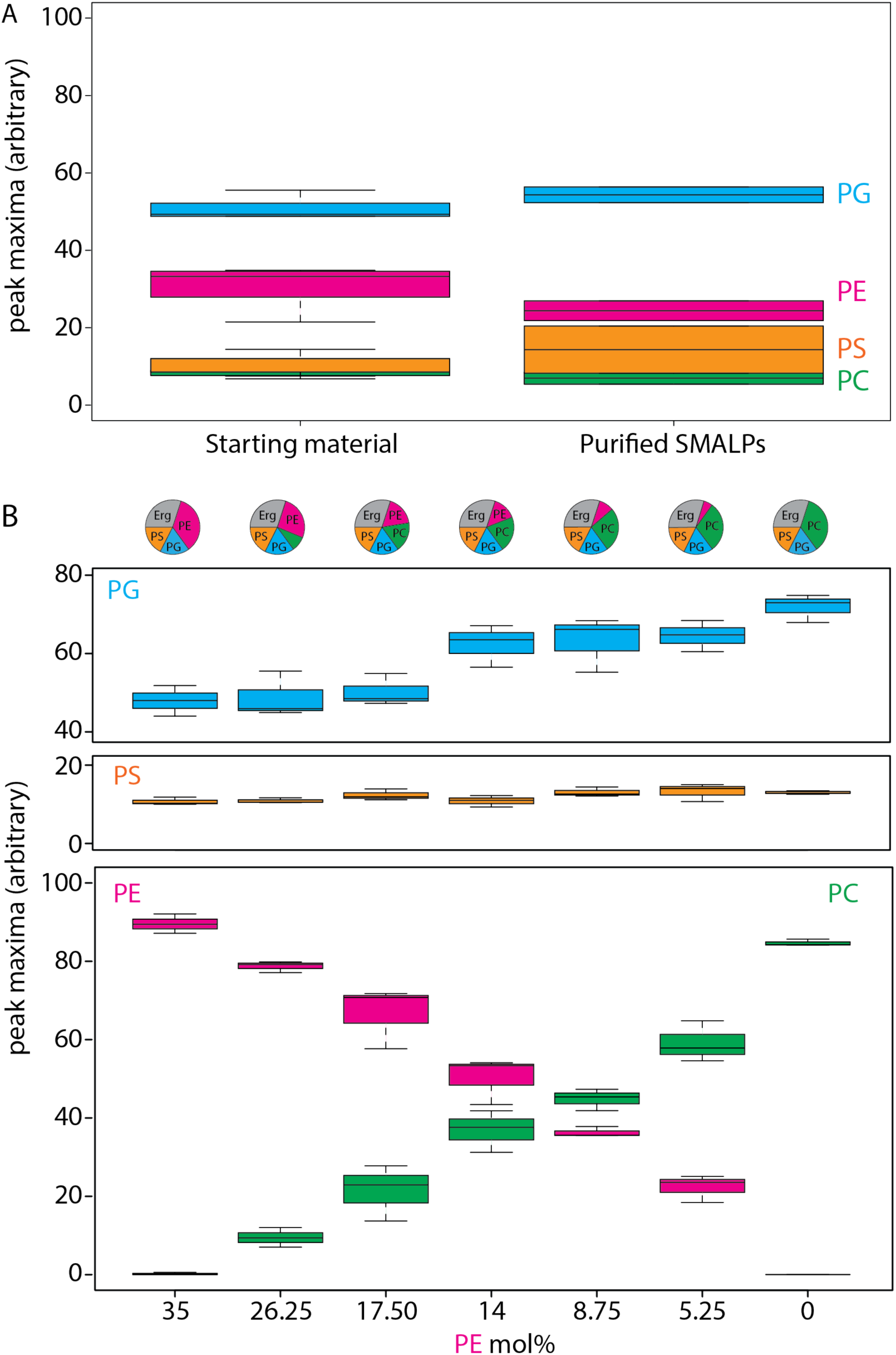
Lipid analysis by mass spectrometry of proteo-liposomes. Boxplot of peak areas of mass spectra corresponding to POPE, POPG, POPS and POPC, present in starting material and purified SMALPs (A) and in respective Lyp1 vesicles (B). Absolute quantities of lipids (mol%) taken for each lipid composition are shown in the pie-graphs above. Two-letter abbreviation and color-coding: PC = Phosphatidyl-Choline (green), PE = Phosphatidyl-Ethanolamine (magenta), PS = Phosphatidyl-Serine (orange) and PG = Phosphatidyl-glycerol (blue). Line within boxplot represents the median. Top and bottom represent the first and third quartile, respectively. Error bars are the minimal and maximal value. Number of experiments = 3 experimental replicates.

**Figure S5:**
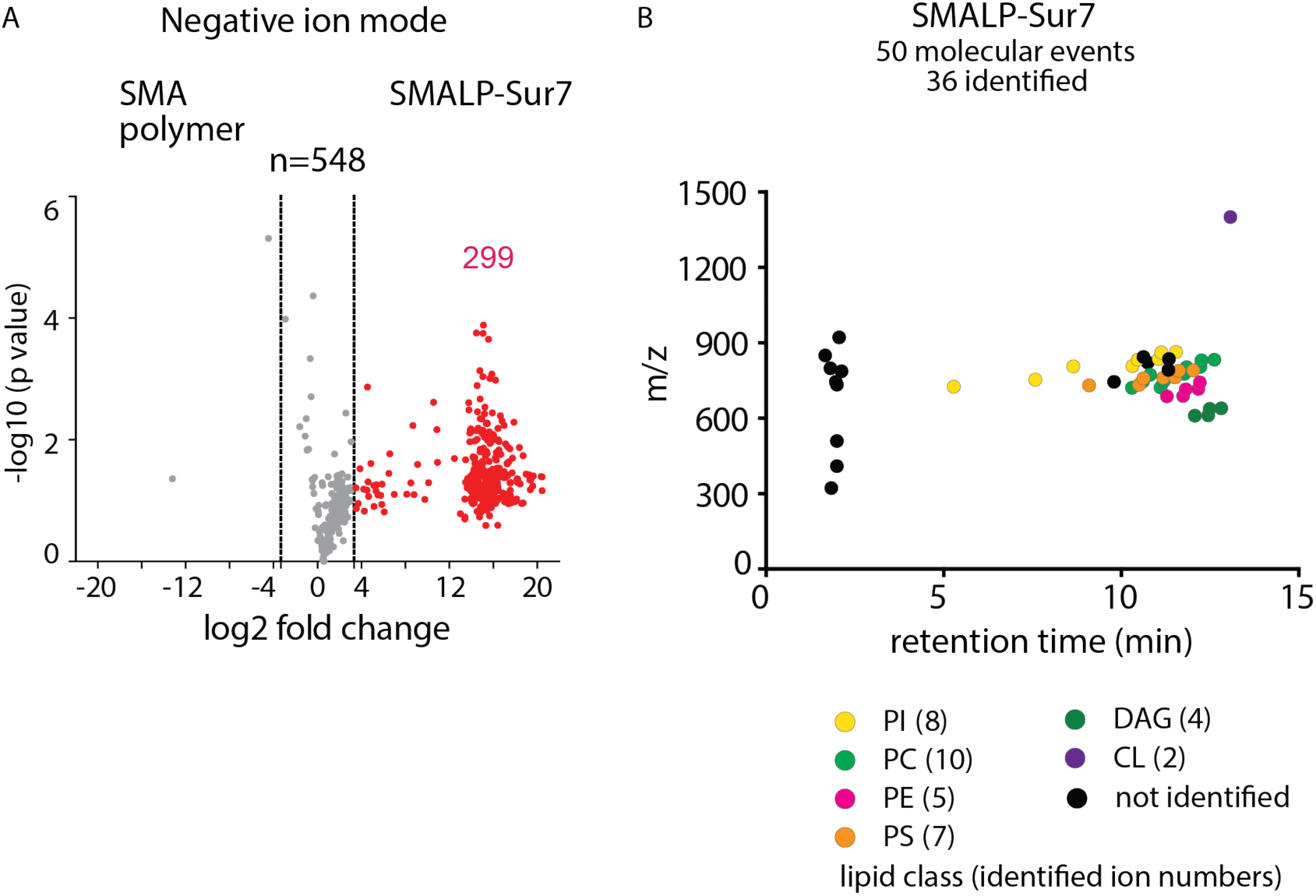
Lipidomics of SMALP-Sur7. (A) Three independently purified samples of SMALP-Sur7 were subjected to lipid extraction, normalization to 1 µM of the input protein and subjected to reversed phase HPLC-QToF-ESI-MS in the negative ion mode. Among 548 ions detected the molecules were considered altered (red) when the signal changed 10-fold. P values were corrected using the Bonferroni method. (B) Using triage methods outlined in Fig. 2, we culled 36 ions with high intensity and lacking redundant detection as isotopes or alternate adducts, whose retention times are shown. Among these, 36 of ions were identified in 7 lipids classes using CID-MS and authentic standards. The number of lipids identified in each class is indicated in the parentheses.

**Figure S6:**
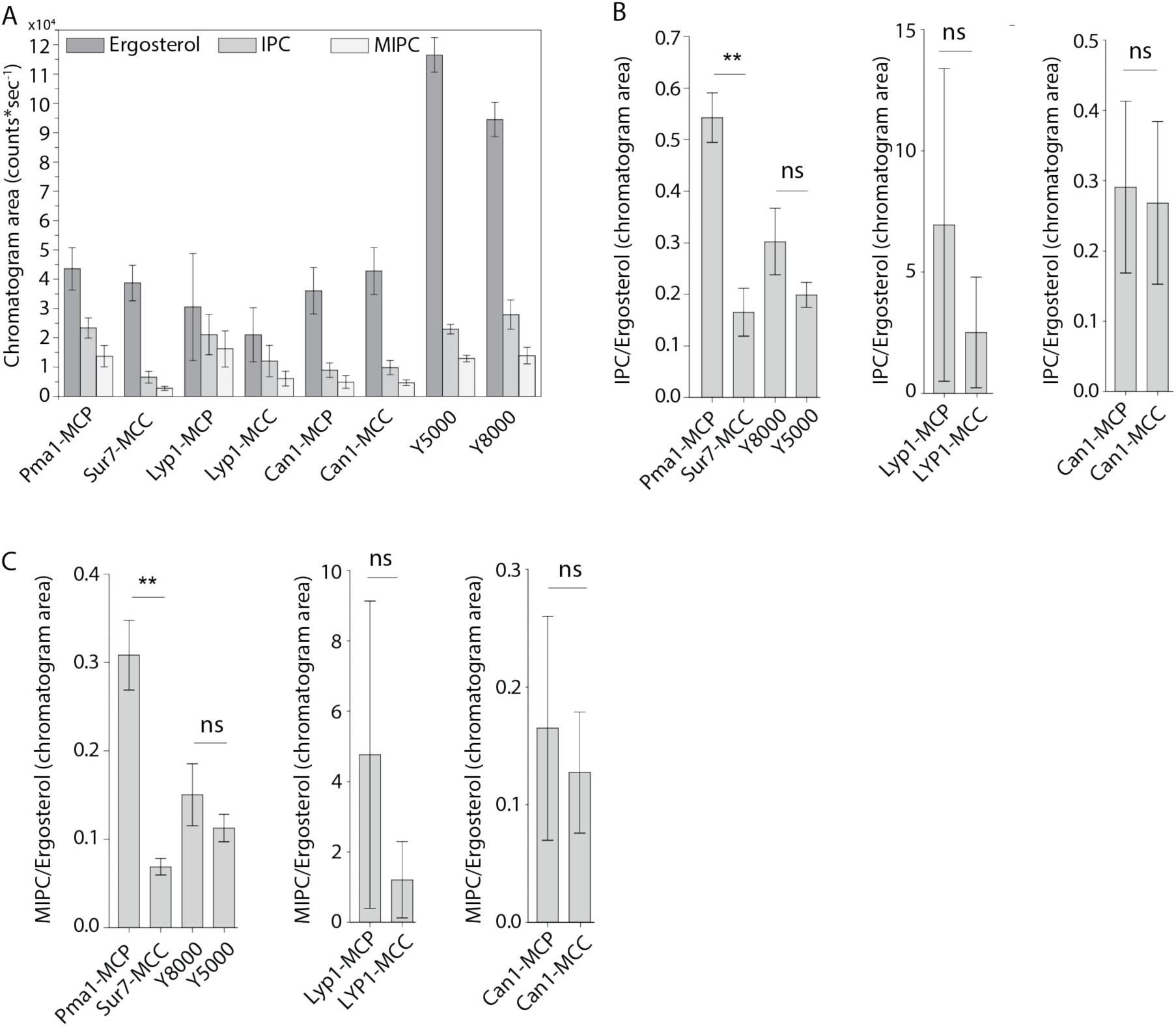
Peak areas and ratios of ergosterol over sphingolipids. (A) Peak areas measured in triplicate with standard error of the mean of the indicated lipid classes are shown according to protein markers that define membrane domains. (B) Ratio of IPC/Ergosterol (C) Ratio of MIPC/Ergosterol. The data are presented as mean +/- standard error. P value was calculated using Student’s t-test. **, P < 0.005, ns = not significant.

**Figure S7:**
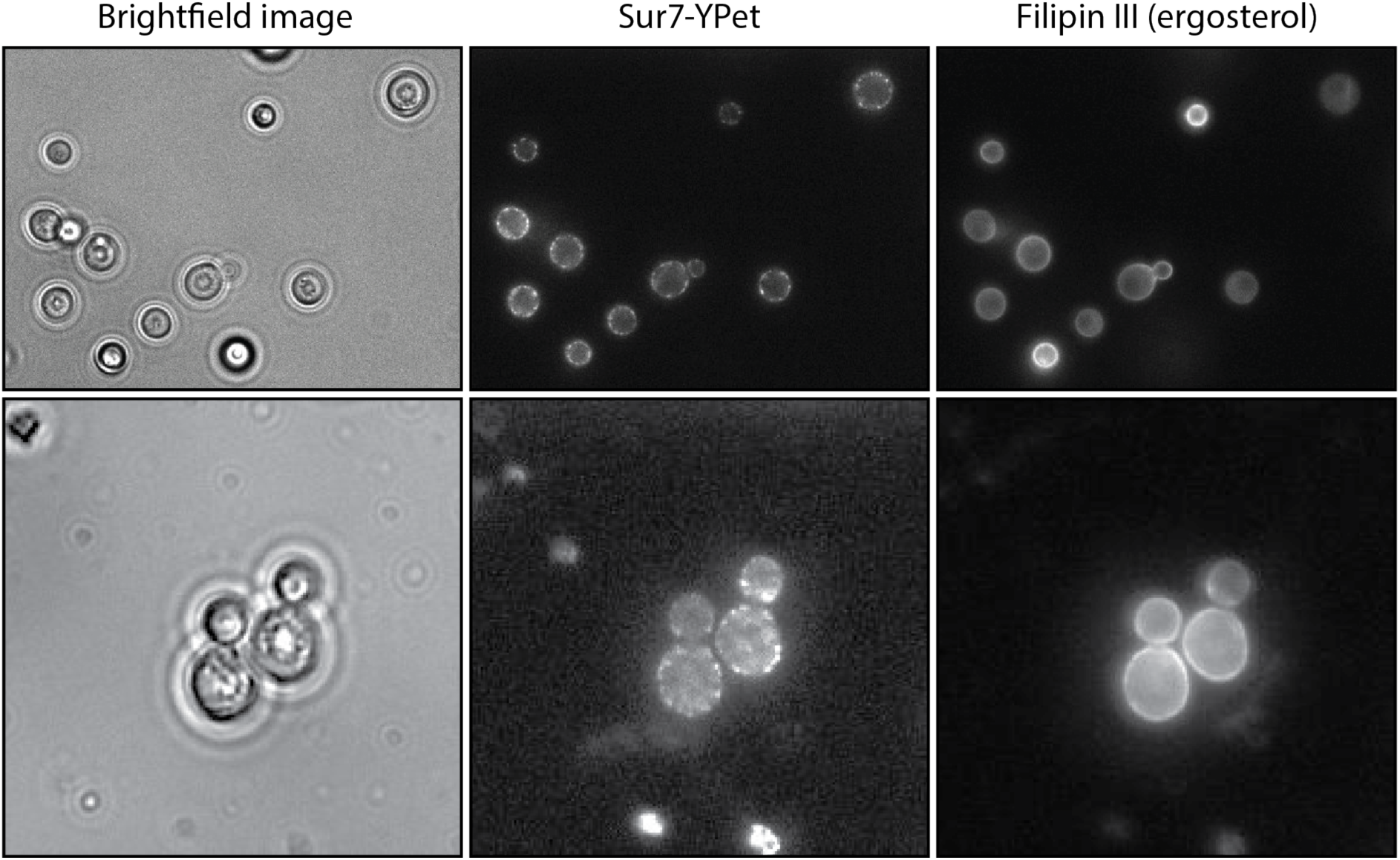
Filipin staining of *S. cerevisiae.* Images of *S. cerevisiae* cells strain Y5007, obtained with visible light (brightfield), showing MCC/eisosomes by 488 nm laser excitation of the MCC marker Sur7-Ypet; ergosterol is visualize by 365 nm excitation of Filipin III.

**Figure S8.**
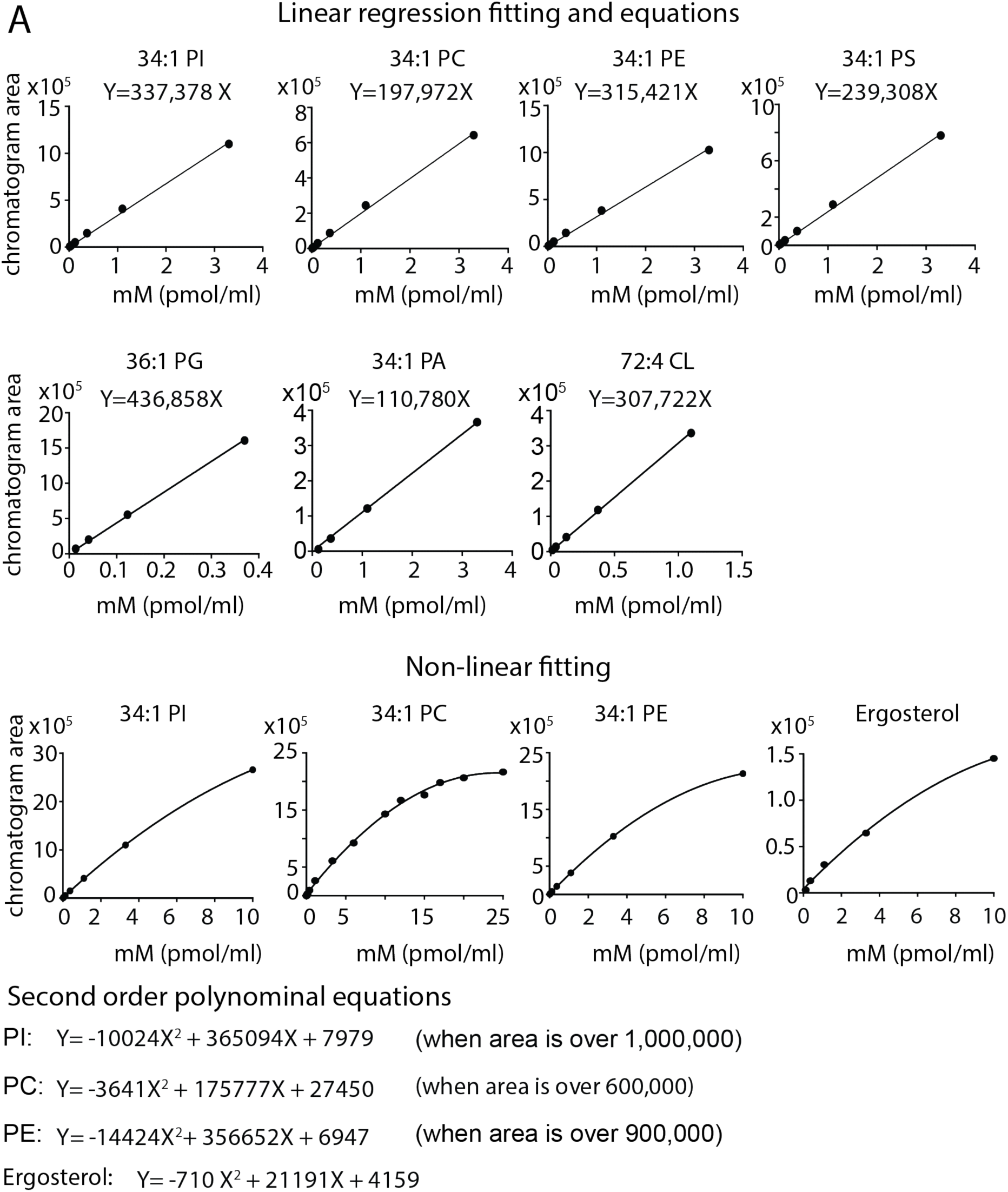
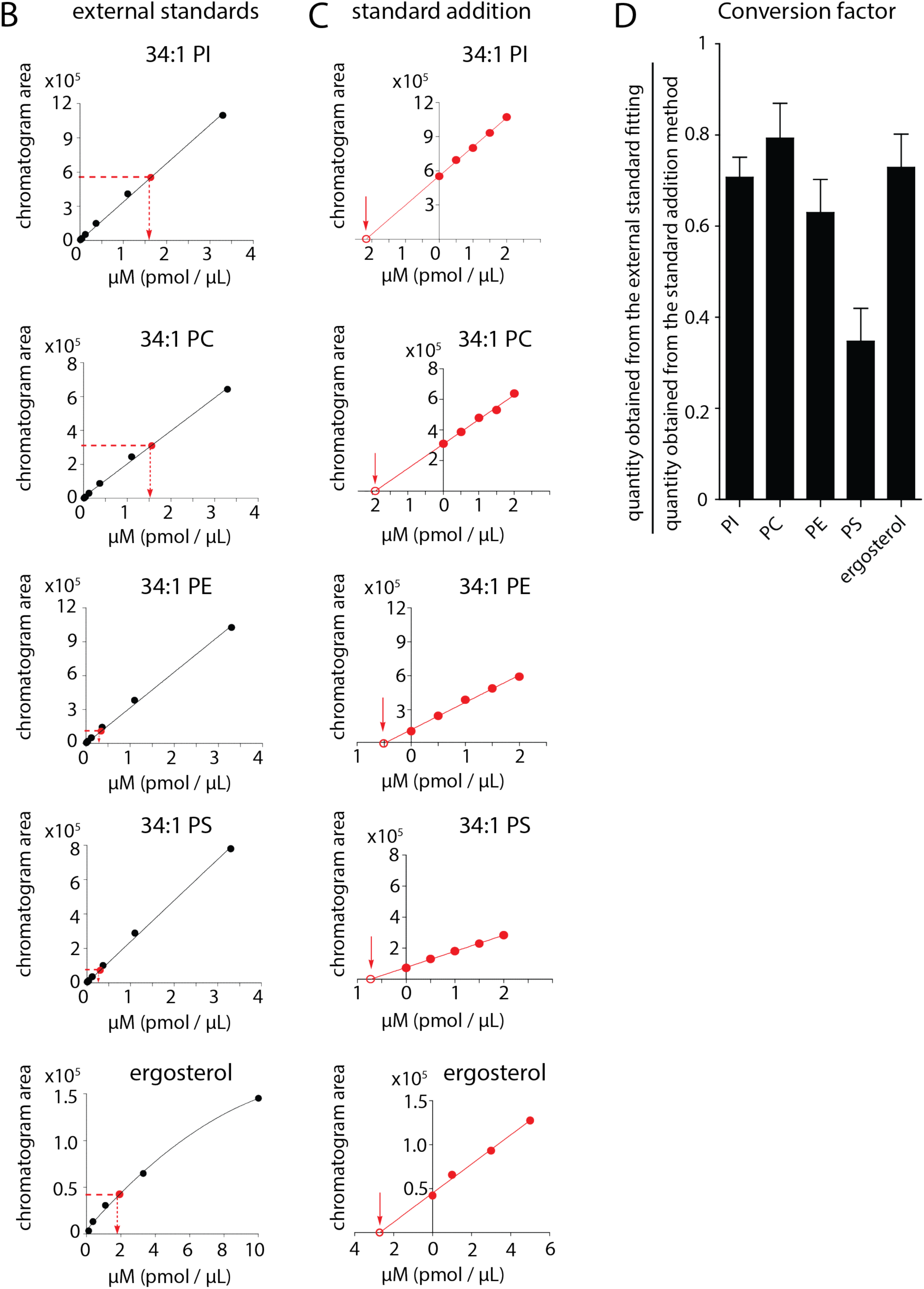
Mass spectrometry-based quantitation using external and internal standards. (A) For each standard, a series of known concentrations were prepared and analyzed by HPLC-MS. The ion chromatogram peak areas from known concentrations were used to generate external standard curves for determining the concentrations of the extracted lipids using linear curve fitting, and non-linear curve fitting for values with zone suppression. (B) Lipid concentrations from SMALPs were estimated using the external standard curve fitting as indicated. (C) Lipid concentrations were also quantified using internal standards by the method of standard addition. Briefly, the lipid extracts were spiked with known concentrations of the synthetic standard, which has the same total fatty acyl chain length and number of unsaturation as compared to the lipid of interest. The ion chromatogram peak areas were plotted against the concentrations of the synthetic standard and extrapolated to the X-axis. (D) The conversion factors were defined as the ratio of the estimated quantity obtained from the external standard fitting to that from the standard addition method. The conversion factors are 0.71, 0.79, 0.63, 0.35, and 0.73 for PI, PC, PE, PS, and ergosterol, respectively. Final mass values are reported according to the estimated amount from the internal standard curve for one lead lipid in each class. The data are presented as mean ± standard deviation of triplicate measurements (SD).

**Figure S9:**
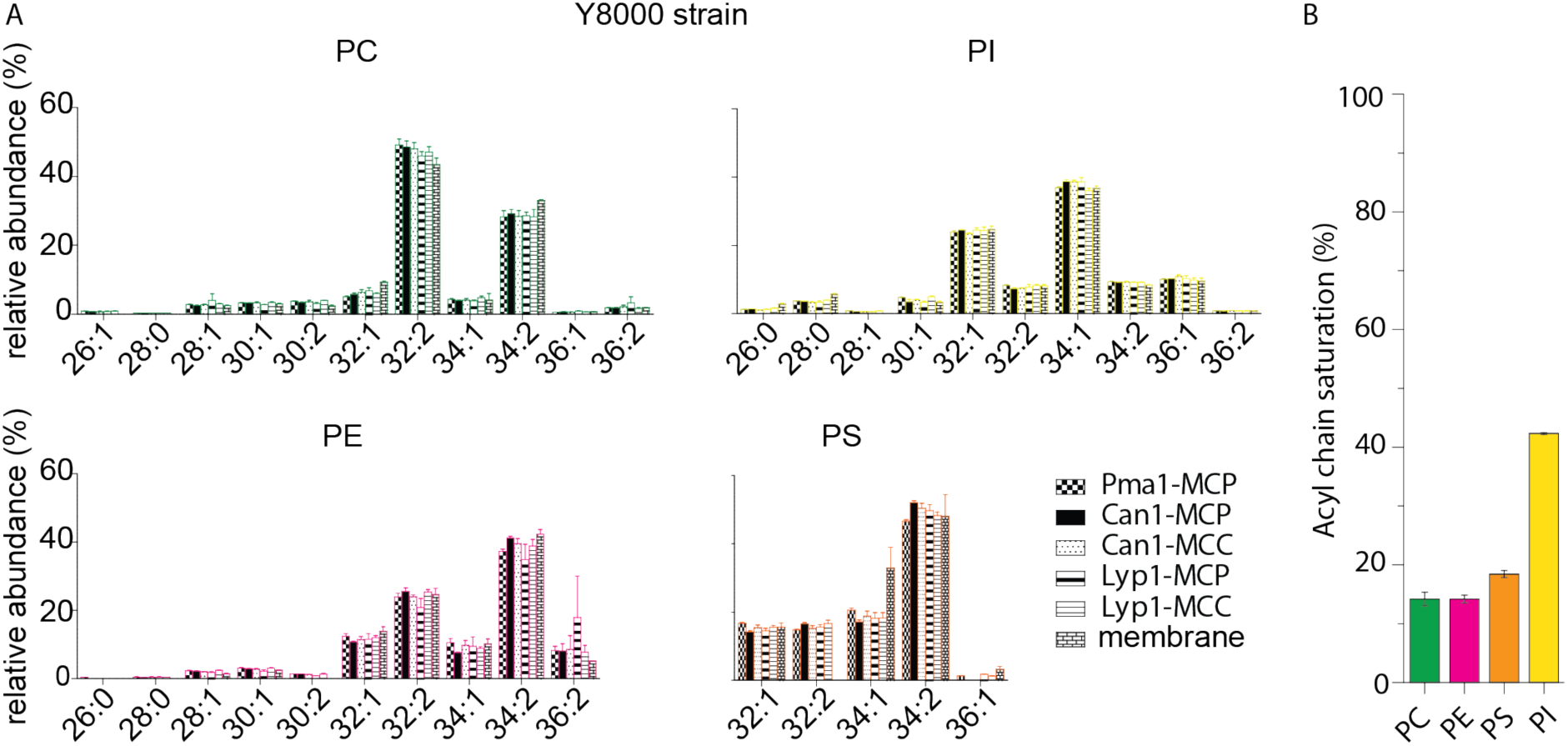
Phospholipid composition of SMALP and membrane samples. (A) Relative abundance of each phospholipid species as a function of acyl chain type for protein-SMALPs. Lyp1-MCC and Can1-MCC refer to SMALPs that have been obtained from proteins trapped in the MCC, using a GFP binding protein fused to the MCC resident Sur7. Lyp1-MCP and Can1-MCP refer to SMALPs that have been obtained from proteins that are not trapped in the MCC and thus correspond to MCP but a minor fraction from the MCC is present. (B) Total acyl chain saturation (%)/phospholipid species.

**Figure S10:**
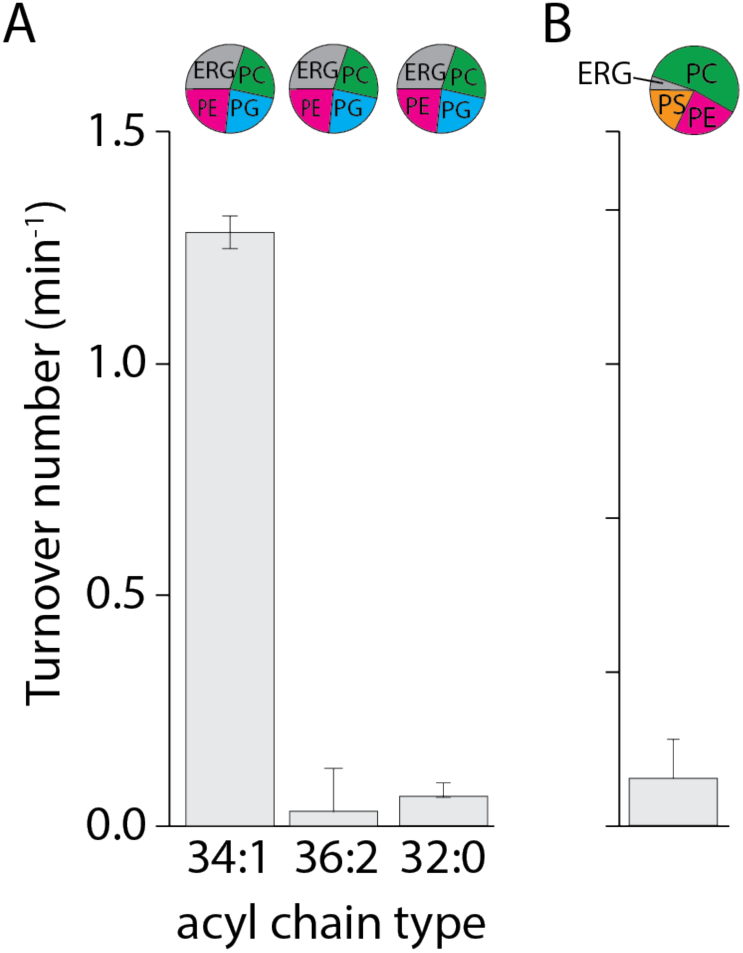
Lyp1 activity as a function of lipid composition. (A) Turnover number of Lyp1p in lipid mixtures with different acyl chains. (B) Turnover number of Lyp1p when measured in a lipid composition based on the lipidomics data from Fig. 3C, using C32:2 PC, C32:2 PE, and C34:1 PS and ergosterol. Triplicate values are shown with bars representing the standard error of the fit of n = 1; variation between replicate experiments is within 20%.

**Figure S11:**
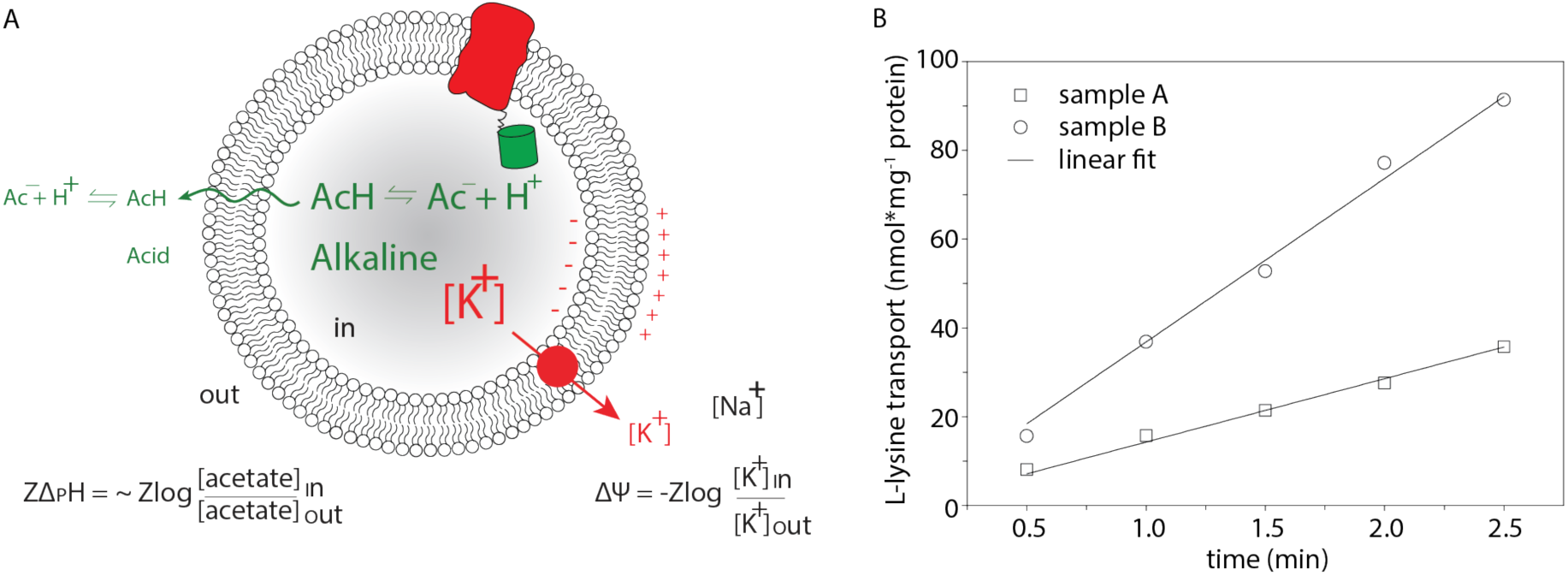
Generation of proton motive force and lysine transport progress curve. A. Schematic showing the generation of a membrane potential (ΔΨ, red arrow) by a valinomycin-mediated potassium diffusion potential and pH gradient (ΔpH, green) formation by an acetate diffusion potential. Together the ΔΨ and ΔpH form the proton motive force (PMF=ΔΨ-ZΔpH, where Z equals 2.3RT/F and R and F are the gas and Faraday constant, respectively, and T is the absolute temperature. B. Transport of lysine by Lyp1-GFP-containing proteoliposomes and data fitting. Lyp1 activity is obtained from the slopes of such lines and converting the rates of transport into turnover numbers

**Figure S12.**
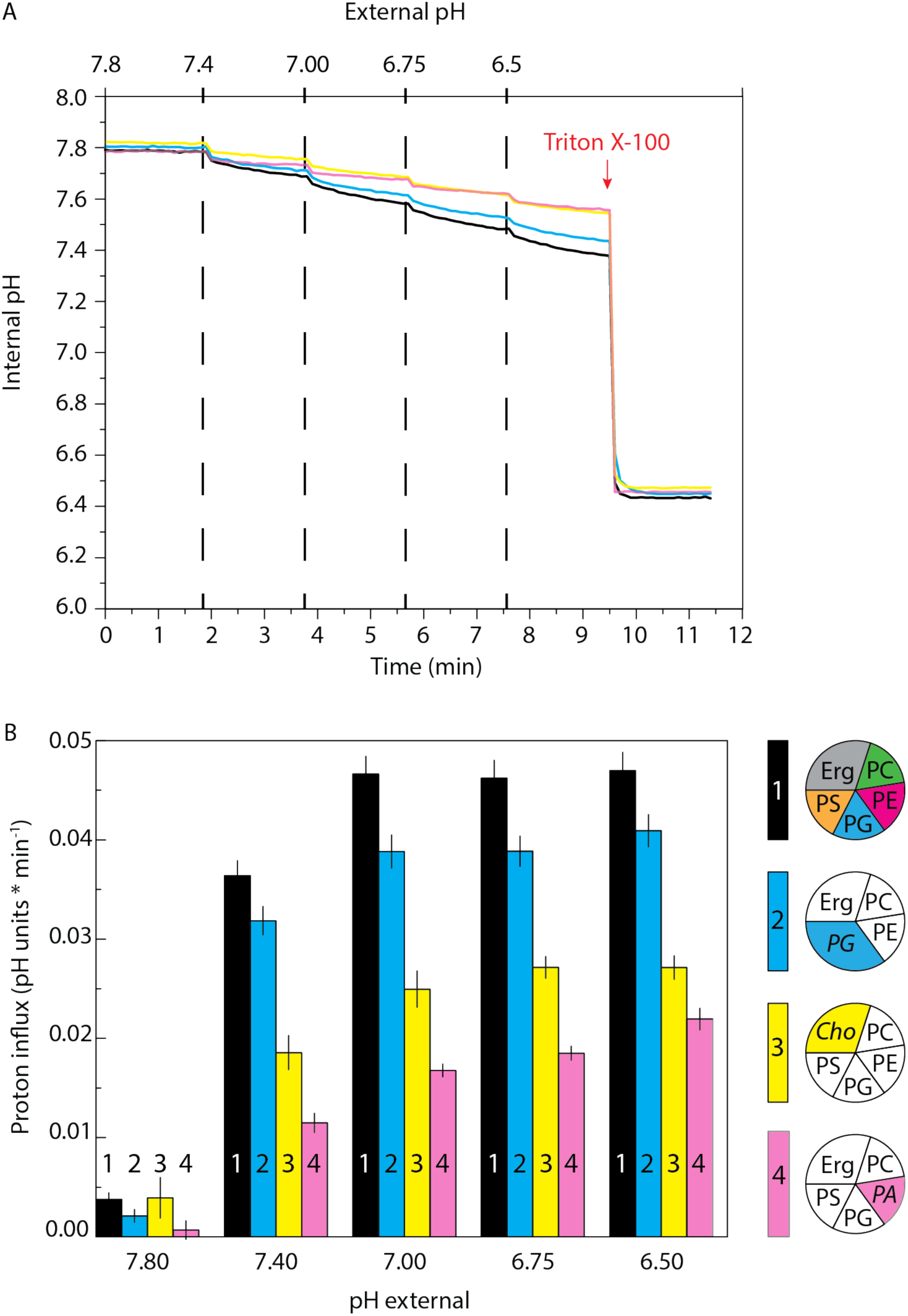
Proton permeability of proteoliposomes. (A) Internal pH is shown as a function of external pH and time for proteoliposomes composed of various lipids. The vesicles were prepared in 10 mM potassium-phosphate pH 7.8. Color code is: POPC/POPE/POPG/POPS/Ergosterol (17,5/17,5/17,5/17.5/30 mol%) = black; POPC/POPE/POPG/Ergosterol (17,5/17,5/35/30 mol%) = Blue; POPC/POPE/POPG/POPS/Cholesterol (17,5/17,5/17,5/17.5/30 mol%) = Yellow; POPC/POPG/POPS/POPA/Ergosterol (17,5/17,5/17,5/17.5/30 mol%) = Pink. The red-arrow indicates Triton X-100 addition, which solubilizes the proteoliposomes and releases pyranine. (B) Proton influx (pH units * min^-1^) is shown as a function of external pH for each lipid composition. The proton influx was calculated from the slope of a linear fit of the data, where dashed lines in Panel A indicate the start and end points used for the fit. Result is representative of three experiments, and error bars are the standard error of the fit.

## References

1. Physical Basis of Self-Organization and Function of Membranes: Physics of Vesicles. Handb Biol Phys [Internet]. 1995 Jan 1 [cited 2019 Sep 10];1:213–304. Available from: https://www.sciencedirect.com/science/article/abs/pii/S1383812106800229?via%3Dihub

2. de la Serna J, Schütz GJ, Eggeling C, Cebecauer M. There Is No Simple Model of the Plasma Membrane Organization. Front Cell Dev Biol. 2016;4:106.

3. Cerbon J, Calderon V. Generation modulation and maintenance of the plasma membrane asymmetric phospholipid composition in yeast cells during growth: their relation to surface potential and membrane protein activity. Biochim Biophys Acta - Biomembr [Internet]. 1995 Apr 12 [cited 2019 Oct 4];1235(1):100–6. Available from: https://www.sciencedirect.com/science/article/pii/000527369400311C?via%3Dihub

4. Solanko LM, Sullivan DP, Sere YY, Szomek M, Lunding A, Solanko KA, et al. Ergosterol is mainly located in the cytoplasmic leaflet of the yeast plasma membrane. Traffic [Internet]. 2018 Mar [cited 2019 Aug 19];19(3):198–214. Available from: http://www.ncbi.nlm.nih.gov/pubmed/29282820

5. Distribution and functions of sterols and sphingolipids. Cold Spring Harb Perspect Biol. 2011;3(5):a004762.

6. Heard GM, Fleet GH. The effects of temperature and pH on the growth of yeast species during the fermentation of grape juice. J Appl Bacteriol. 1988;65(1):23–8.

7. Gray WD. Studies on the Alcohol Tolerance of Yeasts. J Bacteriol [Internet]. 1941 Nov [cited 2019 Oct 22];42(5):561–74. Available from: http://www.ncbi.nlm.nih.gov/pubmed/16560468

8. Bianchi F, Syga Ł, Moiset G, Spakman D, Schavemaker PE, Punter CM, et al. Steric exclusion and protein conformation determine the localization of plasma membrane transporters. Nat Commun. 2018;9(1):501.

9. Gabba M, Frallicciardi J, van’t‘Klooster J, Henderson R, Syga Ł, Mans R, et al. Weak Acid Permeation in Synthetic Lipid Vesicles and Across the Yeast Plasma Membrane. Biophys J [Internet]. 2019 Nov [cited 2019 Dec 4]; Available from: https://linkinghub.elsevier.com/retrieve/pii/S0006349519343413

10. Aresta-Branco F, Cordeiro AM, Marinho SH, Cyrne L, Antunes F, de Almeida RFM. Gel Domains in the Plasma Membrane of Saccharomyces cerevisiae HIGHLY ORDERED, ERGOSTEROL-FREE, AND SPHINGOLIPID-ENRICHED LIPID RAFTS. 2011;286(7):5043–54.

11. Malínská K, Malínský J, Opekarová M, Tanner W. Visualization of Protein Compartmentation within the Plasma Membrane of Living Yeast Cells. Mol Biol Cell. 2003;14(11):4427–36.

12. Malinska K, Malinsky J, Opekarova M, Tanner W. Distribution of Can1p into stable domains reflects lateral protein segregation within the plasma membrane of living S. cerevisiae cells. J Cell Sci. 2004;117(25):6031–41.

13. Gournas C, Gkionis S, Carquin M, Twyffels L, Tyteca D, André B. Conformation-dependent partitioning of yeast nutrient transporters into starvation-protective membrane domains. Proc Natl Acad Sci. 2018;115(14):201719462.

14. Spira F, Mueller NS, Beck G, von Olshausen P, Beig J, Wedlich-Söldner R. Patchwork organization of the yeast plasma membrane into numerous coexisting domains. Nat Cell Biol [Internet]. 2012;14(6):640. Available from: https://www.nature.com/articles/ncb2487

15. Walther TC, Brickner JH, Aguilar PS, Bernales S, Pantoja C, Walter P. Eisosomes mark static sites of endocytosis. Nature. 2006;439(7079):998.

16. Grossmann G, Opekarová M, Malinsky J, Weig-Meckl I, Tanner W. Membrane potential governs lateral segregation of plasma membrane proteins and lipids in yeast. Embo J. 2007;26(1):1–8.

17. Grossmann G, Malinsky J, Stahlschmidt W, Loibl M, Weig-Meckl I, Frommer WB, et al. Plasma membrane microdomains regulate turnover of transport proteins in yeast. J Cell Biol. 2008;183(6):1075–88.

18. Dörr JM, Scheidelaar S, Koorengevel MC, Dominguez JJ, Schäfer M, van Walree CA, et al. The styrene–maleic acid copolymer: a versatile tool in membrane research. Eur Biophys J [Internet]. 2016 Jan 6;45(1):3–21. Available from: http://link.springer.com/10.1007/s00249-015-1093-y

19. Knowles TJ, Finka R, Smith C, Lin Y-P, Dafforn T, Overduin M. Membrane Proteins Solubilized Intact in Lipid Containing Nanoparticles Bounded by Styrene Maleic Acid Copolymer. J Am Chem Soc [Internet]. 2009 Jun 10 [cited 2019 Oct 9];131(22):7484–5. Available from: https://pubs.acs.org/doi/10.1021/ja810046q

20. Itzhak DN, Tyanova S, Cox J, Borner GHH. Global, quantitative and dynamic mapping of protein subcellular localization. Elife. 2016 Jun 9;5(JUN2016).

21. Ghaddar K, Merhi A, Saliba E, Krammer E-M, Prévost M, André B. Substrate-Induced Ubiquitylation and Endocytosis of Yeast Amino Acid Permeases. Mol Cell Biol. 2014;34(24):4447–63.

22. Rothbauer U, Zolghadr K, Muyldermans S, Schepers A, Cardoso MC, Leonhardt H. A versatile nanotrap for biochemical and functional studies with fluorescent fusion proteins. Mol Cell Proteomics. 2008;7(2):282–9.

23. Pardo JJ, Dörr JM, Iyer A, Cox RC, Scheidelaar S, Koorengevel MC, et al. Solubilization of lipids and lipid phases by the styrene–maleic acid copolymer. Eur Biophys J. 2016;46(1):1–11.

24. Scheidelaar S, Koorengevel MC, Pardo J, Meeldijk JD, Breukink E, Killian AJ. Molecular model for the solubilization of membranes into nanodisks by styrene maleic Acid copolymers. Biophys J. 2015;108(2):279–90.

25. Bligh EG, Dyer WJ. A rapid method of total lipid extraction and purification. Can J Biochem Physiol. 1959;37(8):911–7.

26. Layre E, Sweet L, Hong S, Madigan CA, Desjardins D, Young DC, et al. A comparative lipidomics platform for chemotaxonomic analysis of mycobacterium tuberculosis. Chem Biol. 2011 Dec 23;18(12):1537–49.

27. Drage MG, Tsai HC, Pecora ND, Cheng TY, Arida AR, Shukla S, et al. Mycobacterium tuberculosis lipoprotein LprG (Rv1411c) binds triacylated glycolipid agonists of Toll-like receptor 2. Nat Struct Mol Biol. 2010 Sep;17(9):1088–95.

28. Huang S, Cheng T-Y, Young DC, Layre E, Madigan CA, Shires J, et al. Discovery of deoxyceramides and diacylglycerols as CD1b scaffold lipids among diverse groove-blocking lipids of the human CD1 system. Proc Natl Acad Sci [Internet]. 2011 Nov 29 [cited 2019 Sep 10];108(48):19335–40. Available from: http://www.ncbi.nlm.nih.gov/pubmed/22087000

29. Birkinshaw RW, Pellicci DG, Cheng T-Y, Keller AN, Sandoval-Romero M, Gras S, et al. αβ T cell antigen receptor recognition of CD1a presenting self lipid ligands. Nat Immunol [Internet]. 2015 Mar 2 [cited 2019 Sep 10];16(3):258–66. Available from: http://www.nature.com/articles/ni.3098

30. Wun KS, Reijneveld JF, Cheng T-Y, Ladell K, Uldrich AP, Le Nours J, et al. T cell autoreactivity directed toward CD1c itself rather than toward carried self lipids. Nat Immunol [Internet]. 2018 Apr 12 [cited 2019 Sep 10];19(4):397–406. Available from: http://www.ncbi.nlm.nih.gov/pubmed/29531339

31. Bianchi F, van Klooster JS, Ruiz SJ, Luck K, Pols T, Urbatsch IL, et al. Asymmetry in inward- and outward-affinity constant of transport explain unidirectional lysine flux in Saccharomyces cerevisiae. Sci Rep-uk. 2016;6(1):31443.

32. Van Den Brink-Van Der Laan E, Antoinette Killian J, De Kruijff B. Nonbilayer lipids affect peripheral and integral membrane proteins via changes in the lateral pressure profile. Vol. 1666, Biochimica et Biophysica Acta - Biomembranes. 2004. p. 275–88.

33. van der Rest ME, Kamminga a H, Nakano a, Anraku Y, Poolman B, Konings WN. The plasma membrane of Saccharomyces cerevisiae: structure, function, and biogenesis. Microbiol Rev. 1995;59(2):304–22.

34. Greenberg ML, Axelrod D. Anomalously slow mobility of fluorescent lipid probes in the plasma membrane of the yeast Saccharomyces cerevisiae. J Membr Biol. 1993 Jan;131(2):115–27.

35. Krishnamurthy H, Gouaux E. X-ray structures of LeuT in substrate-free outward-open and apo inward-open states. Nature. 2012;481(7382):469.

36. Lomize MA, Pogozheva ID, Joo H, Mosberg HI, Lomize AL. OPM database and PPM web server: Resources for positioning of proteins in membranes. Nucleic Acids Res. 2012 Jan;40(D1).

37. Duschl C, Boncheva M, Vogel H. A miniaturized monolayer trough with variable surface area in the square-millimeter range. Biochim Biophys Acta - Biomembr [Internet]. 1998 May [cited 2019 Nov 18];1371(2):345–50. Available from: https://linkinghub.elsevier.com/retrieve/pii/S0005273698000364

38. Monje-Galvan V, Klauda JB. Modeling Yeast Organelle Membranes and How Lipid Diversity Influences Bilayer Properties. Biochemistry [Internet]. 2015 Nov 17 [cited 2019 Oct 16];54(45):6852–61. Available from: http://www.ncbi.nlm.nih.gov/pubmed/26497753

39. Zinser E, Sperka-Gottlieb CD, Fasch E V, Kohlwein SD, Paltauf F, Daum G. Phospholipid synthesis and lipid composition of subcellular membranes in the unicellular eukaryote Saccharomyces cerevisiae. 1991;173(6):2026–34.

40. Zinser E, Daum G. Isolation and biochemical characterization of organelles from the yeast, Saccharomyces cerevisiae. Yeast. 1995;11(6):493–536.

41. Ejsing CS, Sampaio JL, Surendranath V, Duchoslav E, Ekroos K, Klemm RW, et al. Global analysis of the yeast lipidome by quantitative shotgun mass spectrometry. Proc Natl Acad Sci USA. 2009;106(7):2136–41.

42. Tuller G, Nemec T, Hrastnik C, Daum G. Lipid composition of subcellular membranes of an FY1679-derived haploid yeast wild-type strain grown on different carbon sources. Yeast. 1999;15(14):1555–64.

43. Schneiter R, Brügger B, Sandhoff R, Zellnig G, Leber A, Lampl M, et al. Electrospray ionization tandem mass spectrometry (ESI-MS/MS) analysis of the lipid molecular species composition of yeast subcellular membranes reveals acyl chain-based sorting/remodeling of distinct molecular species en route to the plasma membrane. J Cell Biol. 1999;146(4):741–54.

44. Daum G, Tuller G, Nemec T, Hrastnik C, Balliano G, Cattel L, et al. Systematic analysis of yeast strains with possible defects in lipid metabolism. Yeast. 1999;15(7):601–14.

45. Moreira KE, Schuck S, Schrul B, Fröhlich F, Moseley JB, Walther TC, et al. Seg1 controls eisosome assembly and shape. J Cell Biol. 2012;198(3):405–20.

46. Kraft ML. Sphingolipid Organization in the Plasma Membrane and the Mechanisms That Influence It. 2017;4:154.

47. Bogdanov M, Sun J, Kaback HR, Dowhan W. A Phospholipid Acts as a Chaperone in Assembly of a Membrane Transport Protein. J Biol Chem [Internet]. 1996 May 17;271(20):11615–8. Available from: http://www.jbc.org/lookup/doi/10.1074/jbc.271.20.11615

48. Veld GI, Driessen AJM, Op den Kamp JAF, Konings WN. Hydrophobic membrane thickness and lipid-protein interactions of the leucine transport system of Lactococcus lactis. Biochim Biophys Acta - Biomembr [Internet]. 1991 Jun 18 [cited 2019 Oct 8];1065(2):203–12. Available from: https://www.sciencedirect.com/science/article/pii/000527369190231V?via%3Dihub

49. Karasawa A, Swier LJYM, Stuart MCA, Brouwers J, Helms B, Poolman B. Physicochemical factors controlling the activity and energy coupling of an ionic strength-gated ATP-binding cassette (ABC) transporter. J Biol Chem. 2013;288(41):29862–71.

50. Teo ACK, Lee SC, Pollock NL, Stroud Z, Hall S, Thakker A, et al. Analysis of SMALP co-extracted phospholipids shows distinct membrane environments for three classes of bacterial membrane protein. Sci Rep. 2019;9(1):1813.

51. Laganowsky A, Reading E, Allison TM, Ulmschneider MB, Degiacomi MT, Baldwin AJ, et al. Membrane proteins bind lipids selectively to modulate their structure and function. Nature. 2014;510(7503):172–5.

52. Arora A, Raghuraman H, Chattopadhyay A. Influence of cholesterol and ergosterol on membrane dynamics: a fluorescence approach. Biochem Biophys Res Commun. 2004;318(4):920–6.

53. Kowalczyk L, Ratera M, Paladino A, Bartoccioni P, Errasti-Murugarren E, Valencia E, et al. Molecular basis of substrate-induced permeation by an amino acid antiporter. Proc Natl Acad Sci. 2011;108(10):3935–40.

54. Ghaddar K, Krammer EM, Mihajlovic N, Brohée S, André B, Prévost M. Converting the yeast arginine Can1 permease to a lysine permease. J Biol Chem. 2014;289(10):7232–46.

55. Gournas C, Saliba E, Krammer E-M, Barthelemy C, Prévost M, André B. Transition of yeast Can1 transporter to the inward-facing state unveils an α-arrestin target sequence promoting its ubiquitylation and endocytosis. Mol Biol Cell. 2017;28:2819–32.

56. Schiestl RH, Gietz RD. High efficiency transformation of intact yeast cells using single stranded nucleic acids as a carrier. Curr Genet. 1989;

57. Kaiser C, Michaelis S, Mitchell A, Cold Spring Harbor Laboratory. Methods in yeast genetics : a Cold Spring Harbor Laboratory course manual [Internet]. Cold Spring Harbor Laboratory Press; 1994 [cited 2019 Oct 4]. 234 p. Available from: https://books.google.nl/books/about/Methods_in_yeast_genetics.html?id=u4MMAQAAMAAJ&redir_esc=y

58. Lee SC, Knowles TJ, Postis VLG, Jamshad M, Parslow RA, Lin Y-P, et al. A method for detergent-free isolation of membrane proteins in their local lipid environment. Nat Protoc. 2016;11(7):1149–62.

59. Smith CA, Want EJ, O’Maille G, Abagyan R, Siuzdak G. XCMS: Processing mass spectrometry data for metabolite profiling using nonlinear peak alignment, matching, and identification. Anal Chem. 2006 Feb 1;78(3):779–87.

60. Geertsma ER, Nik Mahmood N a B, Schuurman-Wolters GK, Poolman B. Membrane reconstitution of ABC transporters and assays of translocator function. Nat Protoc. 2008;3(2):256–66.

